# A phosphate starvation response gene (*psr*1-like) is present and expressed in *Micromonas pusilla* and other marine algae

**DOI:** 10.1101/484824

**Authors:** Cara L. Fiore, Harriet Alexander, Melissa C. Kido Soule, Elizabeth B. Kujawinski

## Abstract

Phosphorus (P) limits primary production in regions of the surface ocean, and many plankton species exhibit specific physiological responses to P-deficiency. The metabolic response of *Micromonas pusilla*, an ecologically relevant marine photoautotroph, to P-deficiency was investigated using environmental metabolomics and comparative genomics. The concentrations of some intracellular metabolites were elevated in the P-deficient cells (e.g., xanthine, inosine) and genes involved in the associated metabolic pathways shared a predicted conserved amino acid motif in the non-coding regions of each gene. The presence of the conserved motif suggests that these genes may be co-regulated, and the motif may constitute a regulatory element for binding a transcription factor, specifically that of Psr1 (<u>P</u>hosphate <u>s</u>tarvation <u>r</u>esponse,), first described in the alga, *Chlamydomonas reinhardtii*. A putative phosphate starvation response gene (*psr*1-like) was identified in *M. pusilla* with homology to well characterized *psr*1/*phr*1 genes in algae and plants, respectively. This gene appears to be present and expressed in other marine algal taxa (e.g., *Emiliania huxleyi*) in field sites that are chronically phosphorus-limited. Results from the present study have implications for understanding phytoplankton taxon-specific roles in mediating P cycling in the ocean.

## 1. Introduction

Phosphorus (P) is a critical element for life, found in lipid membranes, genetic material, and energy storage compounds. Because most marine microorganisms preferentially take up P as inorganic phosphate (PO_4_^3-^), concentrations of dissolved PO_4_^3-^ are generally below 1 µM in the surface ocean, (Karl 2014) limiting phytoplankton productivity (Tyrell 1999).

Marine phytoplankton respond to chronic P-deficiency by investing resources in P uptake via inorganic and organic compounds (Chung et al. 2003, Lin et al. 2016), remodeling cellular metabolism and structures (Berdalet et al. 1994, Van Mooy et al. 2009, Martin et al. 2011, Shemi et al. 2016, Kujawinski et al. 2017), and storing P (Dyhrman et al. 2012, Martin et al. 2014). Different mechanisms for enacting these responses have evolved in widely distributed and often sympatric phytoplankton taxa. For example, within the diatom, *Phaeodactylum tricornutum*, proteomics indicated broadly depressed metabolic activity as a response to P-starvation, with down-regulated energy metabolism, amino acid metabolism, nucleic acid metabolism, and photosynthesis, and up-regulated protein degradation, lipid accumulation, and photorespiration (Feng et al. 2016). In contrast, the prymnesiophytes *Prymnesium parvum* and *Emiliania huxleyi,* and the dinoflagellates *Prorocentrum donghaiense* and *Amphidinium carterae*, maintained energy-generating processes (i.e., photosynthesis) and carbon fixation in response to P-deficiency (Lai et al. 2015, Li et al. 2016, Rokitta et al. 2016, Shi et al. 2017). Interestingly, *P. donghaiense* increased nitrate assimilation under P-deficiency (Shi et al. 2017), whereas *E. huxleyi* reduced nitrate uptake with a tight coupling between P and nitrogen (N) pools (Rokitta et al. 2016). These varied strategies all minimize non-critical P utilization and maximize P uptake, and thus play important roles in structuring phytoplankton assemblages in the oceans (Dyhrman et al. 2006, Dyhrman et al. 2009, Martin et al. 2011, Rokitta et al. 2016, Guo et al. 2018).

The cosmopolitan picoeukaryotic groups *Micromonas* and *Ostreococcus* spp. are members of the “green” lineage (Chlorophyta), a monophyletic group including land plants, and are important marine primary producers (Li 1994). Recent culture experiments with *Micromonas pusilla* (Hoppe et al. 2018) and *M. pusilla* Lac38 (Maat et al. 2014) indicated that these strains are likely to be successful under conditions of increased acidification and low nutrient (P) availability. This success may in part, be due to higher efficiency of carbon sequestration for growth under high *p*CO_2_ conditions and/or the ability to remodel photosystem proteins under nutrient-limited conditions (Maat *et al*., 2014; Hoppe *et al*., 2018, Guo *et al*., 2018). Additionally, similar to other phytoplankton (Chung et al. 2003), *M. pusilla* may shift lipid composition to increase non-P containing lipids (Maat et al. 2016, Guo et al. 2018), and up-regulate genes and proteins for P-transporters (Whitney & Lomas 2016, Guo et al. 2018) and genes for polyphosphate accumulation (Bachy et al. 2018). However, some of the broader physiological response mechanisms of *M. pusilla* and other prasinophytes to P-deficiency have yet to be fully explored.

The Chlorophyta share common ancestry with land plants, stemming from a single endosymbiotic event, in contrast to more distantly related phytoplankton groups such as diatoms and haptophytes, which are hypothesized to have undergone multiple endosymbioses (Falkowski et al. 2004; Lewis & McCourt 2004). Consequently, their physiological response to nutritional stressors, like P-deficiency, may share more traits with land plants than with other algal taxa. Indeed, P-deficiency is common in lakes and terrestrial systems (Elser et al. 2007). The physiological response of model chlorophytes such as the mustard plant, *Arabidopsis thaliana*, and the freshwater green alga, *Chlamydomonas reinhardtii*, to P-deficiency led to the identification of a phosphate starvation response gene described as *psr*1 in *C. reinhardtii* (Wykoff et al. 2009) and as *phr*1 in plants (Rubio et al. 2001). This gene encodes for a transcription factor (TF), a protein that binds to specific DNA regions to activate or repress transcription of one or more genes. The specific region of DNA to which TFs bind is typically characterized by a short repetitive nucleotide sequence, or motif, found up- or downstream of a given gene or within introns (Barrett et al. 2012; Franco-Zorrilla et al. 2014). In *A. thaliana,* regulatory motifs for the Phr1 protein were more abundant in known P-responsive genes than in the rest of the genome (Müller et al. 2006), linking Phr1 to the regulation of P-responsive genes. Phr1/Psr1 TFs have not been described, to our knowledge, in marine algae.

Here, we use a combined metabolomic and comparative genomic approach to investigate the impact of P-deficiency on the physiology of *Micromonas pusilla* CCMP1545. Many metabolites are produced by metabolic pathways under genetic regulatory control (e.g., de Carvalho & Fernandes 2010, Markou & Nerantzis 2013); thus, our approach complements recent physiological, transcriptional, and proteomic work (Hoppe et al. 2018, Guo et al. 2018) and provides mechanistic insight into the physiological response to P-deficiency and its underlying genetic regulation. We used a targeted metabolomics approach to analyze the suite of intracellular and extracellular molecules produced by *M. pusilla*. In combination, we employed a comparative genetics approach to identify 1) a *psr*1-like gene in *M. pusilla* and other prevalent marine phytoplankton species and 2) a potential DNA-binding site of the Psr1-like protein in putatively P-responsive genes. We found evidence for the expression of *psr*1-like genes in *M. pusilla* and other marine algae under P-deficient conditions in cultures and in the field. These results have implications for better understanding and predicting the metabolic response of diverse phytoplankton groups to P-deficiency.

## 2. Materials and Methods

### 2.1 Culture of *Micromonas pusilla* CCMP 1545

All glassware was acid-cleaned and combusted at 450°C for at least 4 h. We grew an axenic culture of *Micromonas pusilla* CCMP1545 from the National Center for Marine Algae and Microbiota (Boothbay, ME, USA) in L1-Si media (https://ncma.bigelow.org/algal-recipes) with 0.2-μm filtered (Omnipore filters; Millipore, MA, USA) autoclaved seawater from Vineyard Sound (MA, USA). We maintained the cultures at 22°C under a 12:12 light:dark regime (84 μmol m^-2^ s^-1^). After acclimation to these culture conditions, we split the culture into two parallel cultures of L1-Si media amended with (a) 36 μM phosphate (P-replete) or (b) 0.4 μM phosphate (P-deficient). We maintained these parallel cultures for three generations prior to this experiment (Fig. S1 a,b). At the start of the experiment, we inoculated 9 flasks of each media type with exponential-phase cells to achieve 320 ml and ∼300,000 cells in each flask (33 ml P-replete; 37 ml P-deficient), with three cell-free control flasks. We grew cultures for two weeks (Fig. S2a, b, c) and removed samples approximately 1 hr into the light cycle at experiment initiation (T_0_; day 0), in late exponential growth phase (T_1_; day 4 P-deficient; day 6 P-replete), and in stationary phase (T_2_; day 5 P-deficient; day 13 P-replete).

At each sampling point, we removed 1 ml for cell counts and chlorophyll fluorescence, 30 ml for total organic carbon (TOC; see Methods in the supplement), and 20 ml of filtrate (see below) for nutrients. We monitored cell abundance daily by flow cytometry (Guava, Millipore) and assessed photosystem II efficiency by measuring the variable and maximum chlorophyll fluorescence (F_v_/F_m_) using fluorescence induction and relaxation (FIRe; Satlantic LP, Halifax, Cananda). We used chlorophyll *a* (692 nm) and side scatter of *M. pusilla* cultures to optimize flow cytometry settings. We assessed potential bacterial contamination by viewing DAPI-stained cells at each time point (see extended Methods in the supplement).

### 2.2 Metabolite extraction and instrument methods

Metabolite extraction and analytical methods are optimized for semi-polar compounds with a size range of 100-1000 Da, capturing many primary metabolites such as organic acids, vitamins, and nucleosides. We processed cultures for intracellular and extracellular metabolite extraction as described previously (Fiore et al. 2015, extended Methods in the supplement). For intracellular metabolites, we collected cells via filtration on 0.2 μm PTFE filters (Omnipore, Millipore) and stored them at −80°C. Metabolites were extracted in 500 μl of cold 40:40:20 acetonitrile:methanol:water with 0.1 M formic acid. Extracts were reduced to near dryness under vacuum centrifugation and reconstituted in 500 μl of 95:5 MilliQ water: acetonitrile with deuterated biotin (d2-biotin, final concentration 0.05 μg ml^-1^) added as a high-performance liquid chromatography (HPLC) injection standard; 100 μl of the extract was used for targeted metabolomics analysis. We extracted extracellular metabolites using solid phase extraction (SPE) with 1 g/6 cc PPL cartridges (BondElut, Agilent, Santa Clara, CA, USA) as described previously (Dittmar et al. 2008, Longnecker 2015).

Intracellular and extracellular extracts were used in targeted metabolomics analysis with HPLC (Thermo PAL autosampler and Accela pump) coupled to tandem mass spectrometry (TSQ Vantage, Thermo Fisher Scientific, MA, USA) via a heated electrospray ionization (H-ESI) source operated in both positive and negative ion modes. All 62 standards were analyzed using optimized selected reaction monitoring (SRM) conditions (XCalibur 2.0) in the intracellular samples (Kido Soule et al. 2015), while 23 metabolites with high SPE recovery (Johnson et al. 2017) were analyzed in the extracellular samples. Concentrations of metabolites were calculated based on a five-to-seven point manually-curated external calibration curve (0.5-500 ng ml^-1^). A pooled sample for quality control consisted of 10 parts intracellular extract and one part extracellular extract of each experimental sample and was analyzed every ten injections (see extended Methods in the supplement). A laboratory-fortified pooled sample containing a standard mix was analyzed at the start, middle, and end of the analytical run to assess matrix effects and analytical quality. Chromatographic peaks derived from calibration standards, pooled samples, and experimental samples were manually assessed for quality and metabolite concentrations were exported to Excel. To facilitate comparison with other studies, we also report cell-normalized ratios of the P-deficient to P-replete metabolite concentrations. All metabolomics data are available in MetaboLights (http://www.ebi.ac.uk/metabolights/) with accession number MTBLS295.

While both intracellular and extracellular metabolites were quantified in our experiment, the results and discussion will focus primarily on changes in the intracellular metabolite profile in response to P-deficiency, highlighting compounds that were most relevant to the discovery of the *psr*1-like gene. We further focus the intracellular metabolite comparison on the exponential growth phase (T_1_) to capture clear changes in metabolite profiles between treatments and facilitate comparison to previous studies (Whitney & Lomas, 2016, Bachy et al. 2018). R statistical software (v3.0.2; R Core Team) was used for statistical analyses. We used a Welch’s two sample *t*-test to compare the cell-normalized concentrations of specific metabolites (log transformed to achieve a normal distribution) between P-deficient and P-replete treatments.

### 2.3 Characterization of the phosphate starvation response (psr1) gene and putative DNA-binding motifs

We used the *psr*1 gene sequence from *C. reinhardtii* (Wykoff et al. 1999, NCBI XM_001700501.1) to query the genome of *M. pusilla* using the BLAST option through Integrated Microbial Genomes (IMG, blastx, e-value: 1e^-5^). Conserved domains within putative sequences were characterized using NCBI conserved domain search (CD-search). We then used the putative *M. pusilla psr*1-derived amino acid sequence to query several databases using BLAST (tblastn, e-value: 1e^-5^), including: a re-assembly of the Marine Microbial Eukaryote Transcriptional Sequencing Project (MMETSP) (Johnson *et al*., 2018), NCBI non-redundant protein sequences (nr), the NCBI metagenome proteins (env nr), the *Tara* Oceans eukaryotic unigenes (Carradec, Pelletier et al. 2017), and several eukaryotic algal genomes from IMG (Methods in the supplement). For the *Tara* Oceans data, we selected only the *psr*1-like unigenes from the metatranscriptome assemblies that were assigned the taxonomic classification of *Micromonas*. Within each sample, we normalized the expression of the *psr*1 unigenes to the total abundance of all unigenes taxonomically classified as *Micromonas* (Methods in the supplement). Visualization of the *Tara* Oceans queries were performed using the basemap library in Python and custom Python scripts (see extended Methods in the supplement). For several genes of interest, we compared the distribution of gene expression to the expression of *Micromonas psr*1-like genes in the *Tara* Oceans metatranscriptome using a Kolmogorov-Smirnov test. We tested a relationship between phosphate concentration and normalized expression of *Micromonas psr*1-like genes from *Tara* Oceans metatranscriptomes by comparing the abundance of *Micromonas psr*1-like transcripts above and below the mean concentration of phosphate using a Student’s *t*-test.

We manually checked significant matches for all nucleotide sequences that were collected from database-based (NCBI, IMG, MMETSP, *Tara* Oceans) BLAST searches using the *psr*1-like gene from *M. pusilla*. First, we used conserved domain analysis (Marchler-Bauer et al. 2017) to assess domain similarity to *psr*1 and second, we used the sequences with both characteristic domains of the *C. reinhardtii psr*1-derived amino acid sequence in a multiple sequence alignment to the *C. reinhardtii psr*1-derived amino acid sequence. Sequences that did not align well, and specifically did not align with the myb coiled-coil domain, were removed from further consideration. Sequence alignment and phylogenetic analysis was performed in MEGA 5.2.1 (Tamura et al. 2011) using MUSCLE (Edgar 2004) with default settings in MEGA. Phylogenetic reconstruction of the *psr*1 gene was performed in MEGA using neighbor-joining, maximum parsimony, and maximum likelihood algorithms and bootstrap replication 500 times. For visual clarity, a subset of significant sequence matches was used to create the trees, including representative sequences from each major taxonomic group (see Fig. S4 for the full tree). The tree topology for the different phylogenetic reconstructions was similar for nodes with high bootstrap values (> 50%, data not shown). We present the maximum likelihood tree because this algorithm (Jones-Taylor-Thornton model, default maximum likelihood settings in MEGA) uses the most information in its alignments and typically has high accuracy (Felsenstein 1981, Kishino et al. 1990).

We searched genes of *M. pusilla* CCMP1545 for a regulatory binding element, or motif, where the Psr1-like protein would bind to activate or repress gene transcription. We first searched within a now-archived gene model of *M. pusilla* CCMP1545 (FrozenGeneCatalog_20080206 (ver_1), Table S1), then within the updated gene model from Bachy et al. (2018) (*Micromonas pusilla* CCMP1545/MBARI_models(ver_1), Table 1). Specifically, we analyzed gene sequences of interest for a regulatory element, or motif, using the motif-based sequence analysis tools Multiple Em for Motif Elicitation (MEME Suite) 4.10.1 (Bailey et al. 1994). The genes of interest (Table 1) were selected based on metabolites that were elevated or depressed in concentration in the P-deficient cultures though not necessarily statistically different between treatments (i.e., malate, several amino acids, vitamins, Fig. 1), as well as gene expression information from Bachy et al. (2018). These included genes related to central carbon metabolism, lipid metabolism, vitamin metabolism (pantothenate, folate), and nucleotide metabolism (aspartate). We included genes within the carbohydrate and lipid metabolic pathways based on published work (Hockin et al. 2012, Goncalves et al. 2013) as well as the gene for proline oxidase (POX), a key enzyme up-regulated in *E. huxleyi* in response to N- and P-deficiency (Rokitta et al. 2014, Rokitta et al. 2016). After initial motif discovery, we optimized our putative motif by varying motif width and number within genes of interest and within genes that were not expected to be regulated under P-deficiency, including asparagine synthetase (JGI gene ID: 9681548) and acetyltransferase like/FAD linked oxidase (9687568)). We attempted to identify motifs using the motif comparison tool (Tomtom; Gupta et al. 2007) against the *A. thaliana* database (Protein Binding Microarray (PBM) motifs, (Franco-Zorrilla et al. 2014)) and significant matches were used to guide the optimization of motif discovery settings and inclusive sequences.

**Fig. 1.**
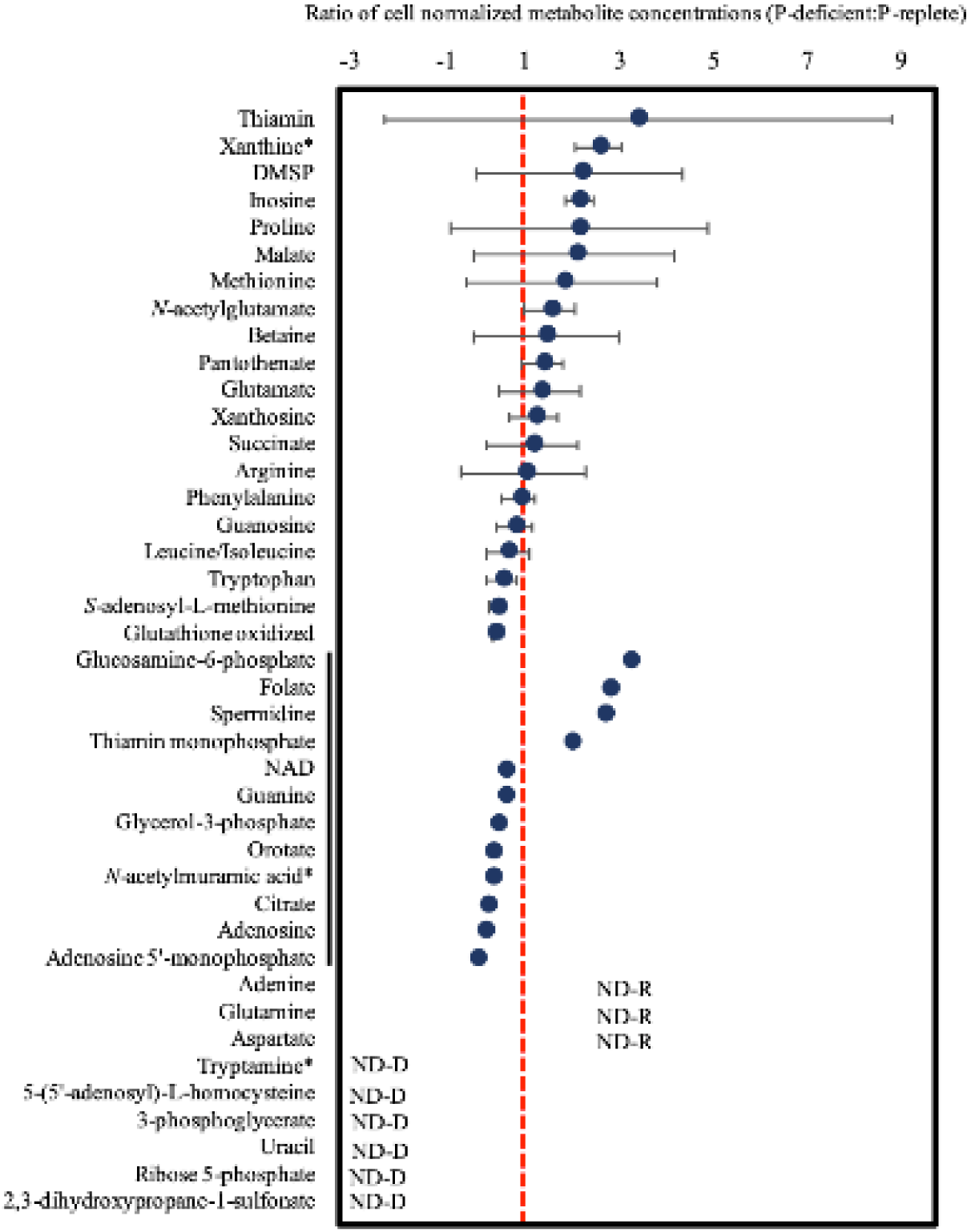
Average ratio of the P-deficient to P-replete intracellular metabolite concentrations during exponential growth (T_1_) in *Micromonas pusilla* CCMP1545. Concentrations are normalized to cell number in each treatment and means with one standard deviation are shown (N=3 unless otherwise noted). Metabolites are listed in order of descending concentration within three groups; those that were detected in enough replicates for standard deviation to be calculated, those where only one ratio could be calculated (black line), and those that were not detected (ND) in any of the three replicates for one treatment (R = P-replete, D = P-deficient). Metabolite names marked with an asterisk indicates a significant difference in concentration between treatments (*t*-test, p < 0.05). Only one replicate in the P-deficient treatment contained a non-zero concentration for *N*-acetylmuramic acid, thus while there was a significant difference in concentration for this metabolite, only one ratio could be calculated. (DMSP = dimethylsulfoniopropionate, NAD = nicotinamide adenine dinucleotide)

**Table 1.** Enzymes that contain a significant motif in the gene sequences of *M. pusilla* CCMP1545 (MBARI_models (ver 1)). Column 2: the Enzyme Commission (E.C.) number or pfam identifier; Column 3: main pathway(s) in which the enzyme is involved; Columns 4-6: fold change and *q*-value of the transcript in P-deficient treatment relative to P-replete conditions are data from Bachy et al. (2018) and corresponding gene ID.

| Enzyme | E.C.<br>Number | Pathway | Log 2 fold change* | *q*-value* | JGI gene<br>ID* |
| --- | --- | --- | --- | --- | --- |
| Glyceraldehyde-3-phosphate dehydrogenase | 1.2.1.12 | Calvin cycle/glycolysis | 2.73 | 0.000037 | 4267 |
| Glyceraldehyde-3-phosphate dehydrogenase | 1.2.1.12 | Calvin cycle/glycolysis | -0.644 | 0.000037 | 700 |
| Phosphoglycerate kinase | 2.7.2.3 | Glycolysis | -0.699 | 0.000037 | 7661 |
| Pyruvate kinase | 2.7.1.40 | Glycolysis | -1.32 | 0.000037 | 9370 |
| Pyruvate dehydrogenase | 1.2.4.1 | TCA<sup>a</sup> cycle/Glycolysis | 0.756 | 0.000037 | 3535 |
| Citrate synthase | 2.3.3.1 | TCA cycle | 1.68 | 0.000037 | 1278 |
| Citrate synthase | 2.3.3.1 | TCA cycle | 1.10 | 0.000037 | 3714 |
| Aconitase | 4.2.1.3 | TCA cycle | 1.00 | 0.000037 | 2901 |
| Succinate dehydrogenase | 1.3.5.1 | TCA cycle and ETC<sup>b</sup> | 1.16 | 0.001515 | 2182 |
| Fumarase | 4.2.1.2 | TCA cycle | 1.68 | 0.000037 | 600 |
| Fumarase | 4.2.1.2 | TCA cycle | 1.73 | 0.000068 | 6662 |
| Copper amine oxidase | 1.4.3.21-22 | Arginine and proline metabolism | 2.87 | 0.000037 | 5863 |
| Aspartate transcarbamylase | 2.1.3.2 | Pyrimidine biosynthesis | -0.609 | 0.000037 | 3615 |
| Nucleoside phosphatase | 3.6.1.3 | Pyrimidine metabolism | 0.711 | 0.000037 | 3322 |
| Pantoate-beta-alanine ligase | 6.3.2.1 | Pantothenate and CoA biosynthesis | -0.616 | 0.000037 | 5304 |
| Lysophospholipase | 3.1.1.5 | Glycerophospholipid metabolism | 1.65 | 0.000037 | 3652 |
| Acyl-CoA synthetase | 6.2.1.3 | Fatty acid metabolism | 3.20 | 0.000037 | 1751 |
| Long-chain acyl-CoA synthetases | 6.2.1.3 | Fatty acid metabolism | 0.736 | 0.000037 | 416 |
| Na<sup>+</sup>/PO<sub>4</sub> transporter | PF02690 | inorganic nutrient transport | 4.54 | 0.000037 | 5963 |

\*From Bachy et al. (2018) <sup>a</sup>Tricarboxylic acid cycle <sup>b</sup>Electron transport chain <sup>c</sup>Triacylglycerol

Using the selected sequences and optimized parameters, we searched for motifs that are overrepresented in the query sequences (“positive”) relative to a set of background sequences (“negative”), the latter defined using the genome scaffolds. We used positive and negative sequences to create a position specific prior (PSP) distribution file (Bailey et al. 1994), for further discriminative motif discovery within *M. pusilla* gene sequences of interest (parameters: minw = 5, maxw = 10, nmotifs = 3, mod = oops (one occurrence per sequence)). Each gene sequence consisted of concatenated 500 nucleotides upstream and downstream of the gene as well as untranslated regions (UTRs) and intronic regions, thereby excluding the coding regions.

Two recent publications provided transcriptomes for testing our hypotheses regarding expected differentially expressed (DE) genes in P-deficient and P-replete conditions. Whitney & Lomas (2016) published transcriptomes of *M. commoda* (formerly *M. pusilla*) RCC299, while Bachy et al. (2018) published transcriptomes of *M. pusilla* CCMP1545, under P-deficient or limiting (respectively) and P-replete conditions. Genes discussed as DE here were defined by each study (Whitney & Lomas 2016, Bachy et al. 2018). We note that the each *Micromonas* experiment used slightly different growth conditions and were sampled at different stages of growth. Thus, we focus our comparisons on work in the same strain (*M. pusilla* CCMP1545, Bachy et al. 2018), with brief complementary discussions about *M. commoda* (see extended Results in the supplement). Bachy et al. (2018) used semi-continuous cultures grown under continuous light and were grown with a moderately higher concentration of P (2.5 μM) than in our experiment (0.4 μM P). Despite these differences, the transcriptional data of *M. pusilla* under P-limitation provided a preliminary test of our hypotheses derived from the metabolomics analyses with *M. pusilla* CCMP1545 and *in silico* genomic analysis. We further reasoned that a signal observed across species and experiments would likely represent a conserved and biologically meaningful response to P-deficiency.

## 3. Results

### 3.1 Metabolic response of Micromonas pusilla CCMP1545 to phosphorus deficiency

Metabolite concentrations were generally lower under P-deficiency in the extracellular (Fig. S3) fraction relative to the P-replete cultures, while there was a mixed response in metabolite concentrations in the intracellular fraction (Fig. 1). The extracellular metabolite concentrations represent cumulative changes in the media over time. In stationary phase, nearly all of the quantified extracellular metabolites were less abundant in the P-deficient samples than the P-replete samples (Fig. S3). In contrast, several of the intracellular metabolites were more abundant during exponential growth (T_1_) under P-deficiency, including nucleobases (xanthine, adenine), the nucleoside inosine, amino acids (glutamine, aspartate), the dicarboxylic acid malate, and vitamins (thiamin, pantothenate, folate) (Fig. 1). None of the extracellular metabolites and only a few intracellular metabolites showed statistically significant differences between treatments, including: (a) the nucleobase xanthine, significantly elevated under P-deficiency (*t*-test, p = 0.01), and (b) the amino acid derivatives, *N*-acetylglutamate and tryptamine, significantly higher in P-replete samples (*t*-test, p = 0.01, p = 0.02, respectively) (Fig. 1). We observed decreases in the intracellular ratios of purine nucleosides (guanosine, xanthosine) to their nucleobases (guanine, xanthine) under P-deficiency at T_1_ (Fig. 2). The decrease in the xanthosine:xanthine ratio was statistically significant (*t*-test, p = 0.03). The change in the guanosine:guanine ratio could not be statistically tested as guanine was below detection in two of three P-replete cultures (Fig. 2). Intracellular malate, succinate, and citrate exhibited divergent responses, although their abundances were not significantly different between treatments (Fig. 1). Malate was more abundant in P-deficient cells, while succinate concentrations were similar between treatments and replicates and citrate was detected in only one of three replicates in each treatment (Fig. 1). We noted similarly varied responses to P-deficiency in the purine nucleosides xanthosine and inosine, with invariant xanthosine abundances but higher average concentrations and variability of inosine (Figs. 1,2).

**Fig. 2.**
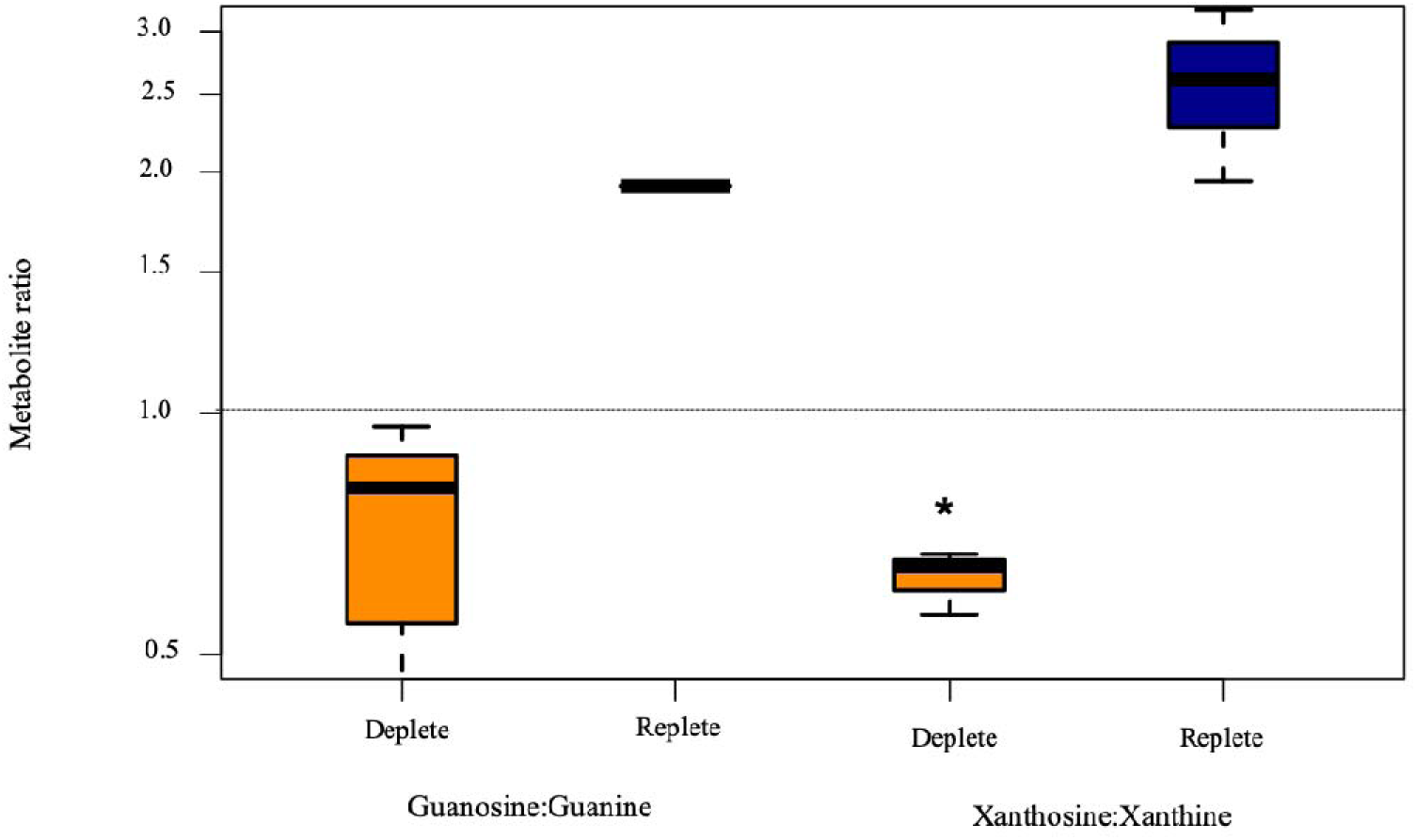
The intracellular ratios of two purine nucleosides to their nucleobases for *Micromonas pusilla* CCMP1545 in P-deficient (orange) and P-replete (blue) treatments during exponential growth (T_1_). Xanthosine and xanthine, and guanosine and guanine were quantified in the targeted metabolomics method and normalized to cell abundance. The average ratio and high and low values are shown based on three replicate cultures. The asterisk marks a significant difference between the treatments (*t*-test, p = 0.03). The dashed line represents a ratio of 1. Two samples in the P-replete treatment had guanine levels below the limit of detection and the one ratio is shown as a line.

Metabolite concentrations are, in part, determined by genetic regulation of the enzymes producing or degrading them and we hypothesized that the varied metabolite responses could provide insight into the regulation of these genes. The description of a phosphate starvation response (*psr*1) gene containing a myb DNA-binding domain (Wykoff et al. 1999), motivated us to investigate the presence of *psr*1 and a myb-like motif in genes that could be regulated by the *psr*1 gene product in *M. pusilla* and other marine algae.

### 3.2 Characterization of the phosphate starvation response (psr1) gene in marine algae

We found four statistically-significant sequences (Table 2) in response to querying the now-archived *M. pusilla* CCMP1545 genome with the *psr*1 gene from *C. reinhardtii* (strain cc125, NCBI accession AF174532). Only one of these sequences (JGI ID 61323) had the two characteristic myb domains of the *psr*1 gene in *C. reinhardtii* (myb-like DNA-binding domain, SHAQKYF class (TIGR01557) and myb predicted coiled-coil type transfactor, LHEQLE motif (pfam PF14379)) (Fig. 3a). The glutamine-rich regions, characteristic of TF activation domains, and putative metal binding domain originally described for *psr*1 in *C. reinhardtii* (Wykoff et al. 1999) were not detected in the putative *M. pusilla psr*1 gene, thus we refer to this as a *psr*1-like gene. Conserved domain analysis (Marchler-Bauer et al. 2017) revealed two other domains in the putative *psr*1-like-derived amino acid sequence from *M. pusilla*: 1) PLN03162 super family (NCBI cl26028), a provisional golden-2 like transcription factor, and 2) a DUF390 super family domain (NCBI cl25642), comprising proteins of unknown function. A comparison of the psr1-like gene in *M. pusilla*, *M. commoda*, *C. reinhardtii*, and *A. thaliana* is provided in Table S2.

**Fig. 3.**
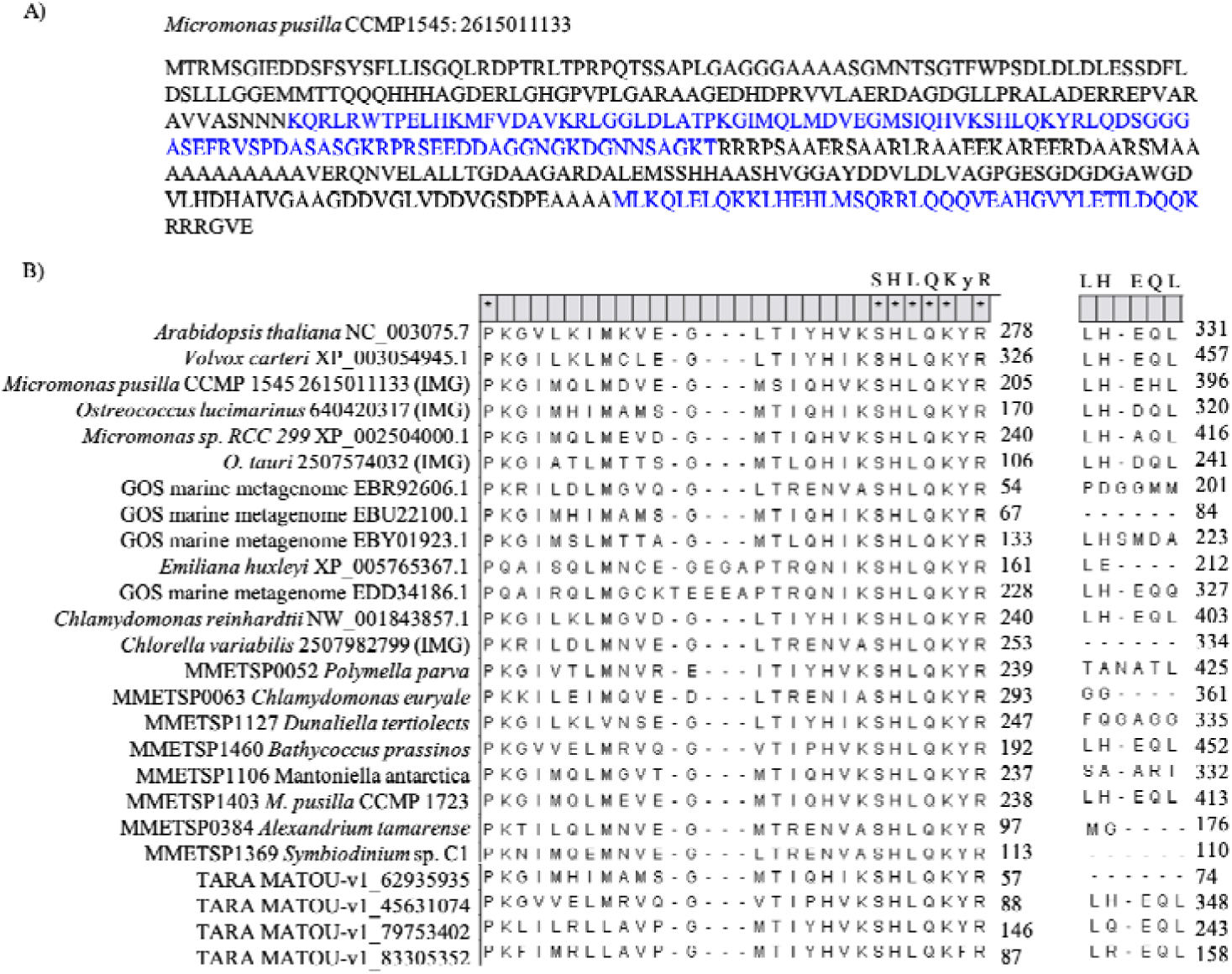
Predicted amino acid sequences of the *psr*1-like gene in *Micromonas pusilla* CCMP1545 and a subset of other organisms. The amino acid sequence (with IMG gene ID) is shown as described in the Joint Genome Institute Integrated Microbial Genomes *M. pusilla* CCMP1545 database. The myb-like DNA-binding and myb coiled-coiled domains are highlighted in blue (A). Predicted amino acid sequence alignment of *psr*1 and *psr*1-like genes in the region of the myb-like DNA-binding domain (SHAQKYF class) and LHEQL coiled-coil domain (B). Asterisks at the top of each column indicate 100% conserved residues across species surveyed. Numbers at the end of each row indicate position of the last shown residue in the amino acid sequence. Accession (MMETSP, NCBI, *Tara*) or gene identification (IMG) numbers are listed for each sequence.

**Table 2.** Gene matches to *psr*1 in *Micromonas pusilla* CCMP1545 as originally discovered in the Joint Genome Institute (JGI) Integrated Microbial Genomes (IMG) database and the corresponding gene in the updated gene model for CCMP1545 (MBARI_models (ver 1)). The *M. pusilla* CCMP 1545 database in IMG contained 10,660 sequences and 4,795,637 letters. The top-scoring gene was considered to be a *psr*1-like gene and the JGI gene ID is shown in parentheses. The corresponding *psr*1-like gene in the updated gene model is shown for reference, and includes the JGI gene ID, the gene description, and the E-value from the BLAST search (see Methods) using the top scoring gene (613233) from CCMP1545.

| IMG Gene ID | Locus tag | Bit score | E-value |
| --- | --- | --- | --- |
| 2615011133 (JGI 61323) | MicpuC2.est_orfs.1_306_4269596:1 | 90.5 | 2 x 10<sup>-19</sup> |
| 2615008475 | MicpuC2.EuGene.0000130210 | 74.3 | 4 x 10<sup>-14</sup> |
| 2615010887 | MicpuC2.est_orfs.10_1861_4270447:1 | 70.5 | 3 x 10<sup>-13</sup> |
| 2615007841 | MicpuC2.EuGene.0000090108 | 67.8 | 4 x 10<sup>-12</sup> |
| JGI ID 360 | Myb domain-containing protein |  | 8 x 10<sup>-156</sup> |

We identified putative *psr*1-like genes or transcripts in cultured isolates as well as field datasets (Figs. 3b, 4, 5, Table S3), representing diverse taxa such as Dinophyta (e.g., the symbiotic clade C *Symbiodinium* sp.), and Haptophyta (e.g., the coccolithophore *E. huxleyi)*. In our phylogenetic analysis (Fig. 4, Fig. S4 for full tree and extended Results in the supplement), representative sequences clustered by taxonomic origin only to some extent. For example, the prasinophytes, including *M. pusilla* CCMP1545, generally clustered together, as did the haptophytes, including *E. huxleyi*. However, other clusters included a mix of taxonomic groups such as the dinoflagellate *Alexandrium tamarense*, the glaucophyte *Gloeochaete wittrockiana*, and the green alga *Chlorella vulgaris* (Fig. 4).

**Fig. 4.**
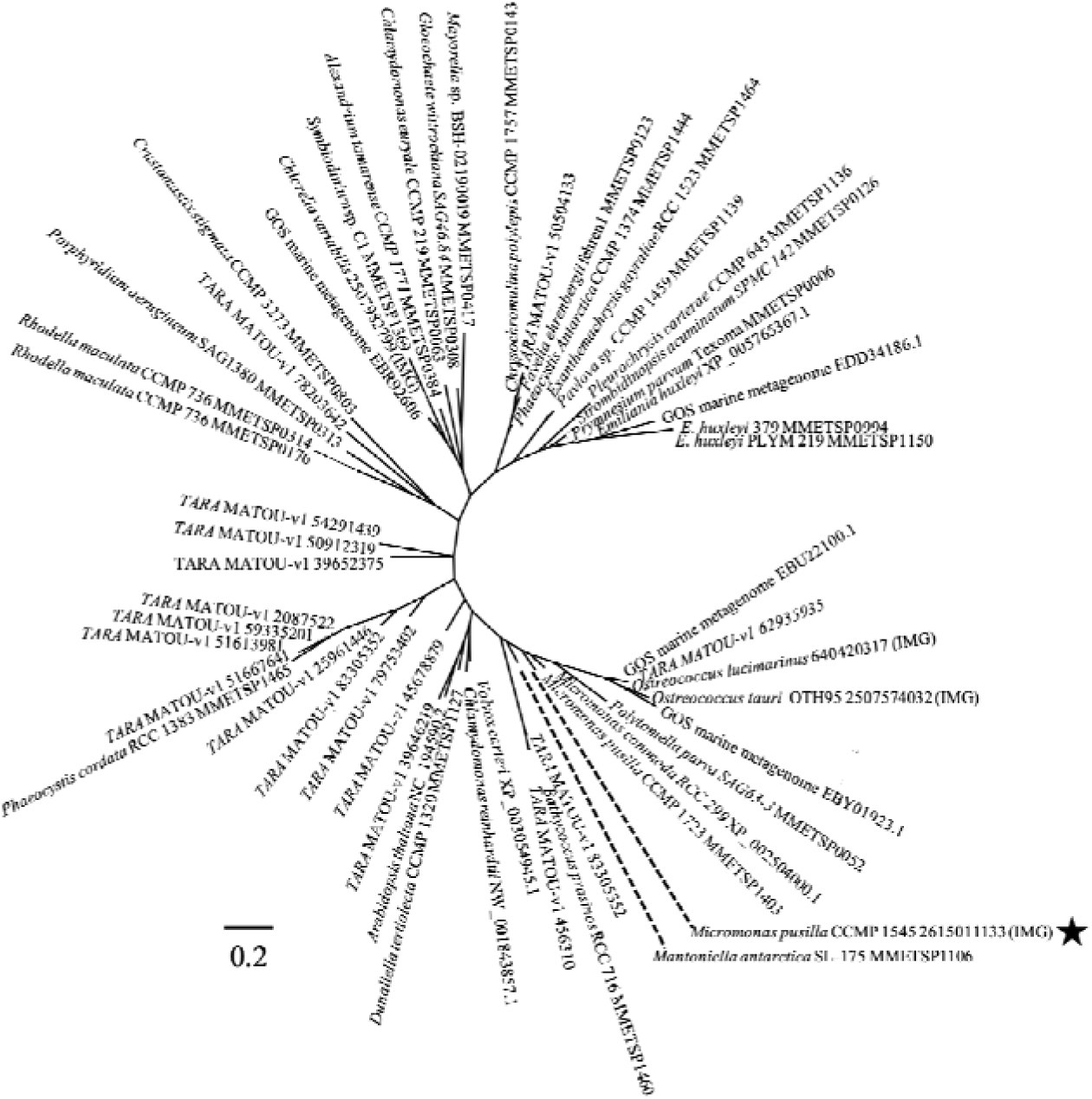
Occurrence and phylogenetic relationship of *psr*1-like genes in marine phytoplankton. Maximum likelihood method based on the JTT matrix-based model (Jones *et al*., 1992). The tree with the highest log likelihood (−2442.94) is shown and is based on the derived amino acid sequences for *psr*1 and *psr*1-like genes from eukaryotic phytoplankton. The analysis involved 55 amino acid sequences. All positions containing gaps and missing data were eliminated yielding a total of 43 positions in the final dataset. Evolutionary analyses were conducted in MEGA7 (Kumar *et al*., 2016) Accession or IMG gene identification numbers are shown with the taxon name and the star indicates *Micromonas pusilla* CCMP1545 cultured in this study. The tree is unrooted and the scale indicates the number of amino acid substitutions per site.

**Fig. 5.**
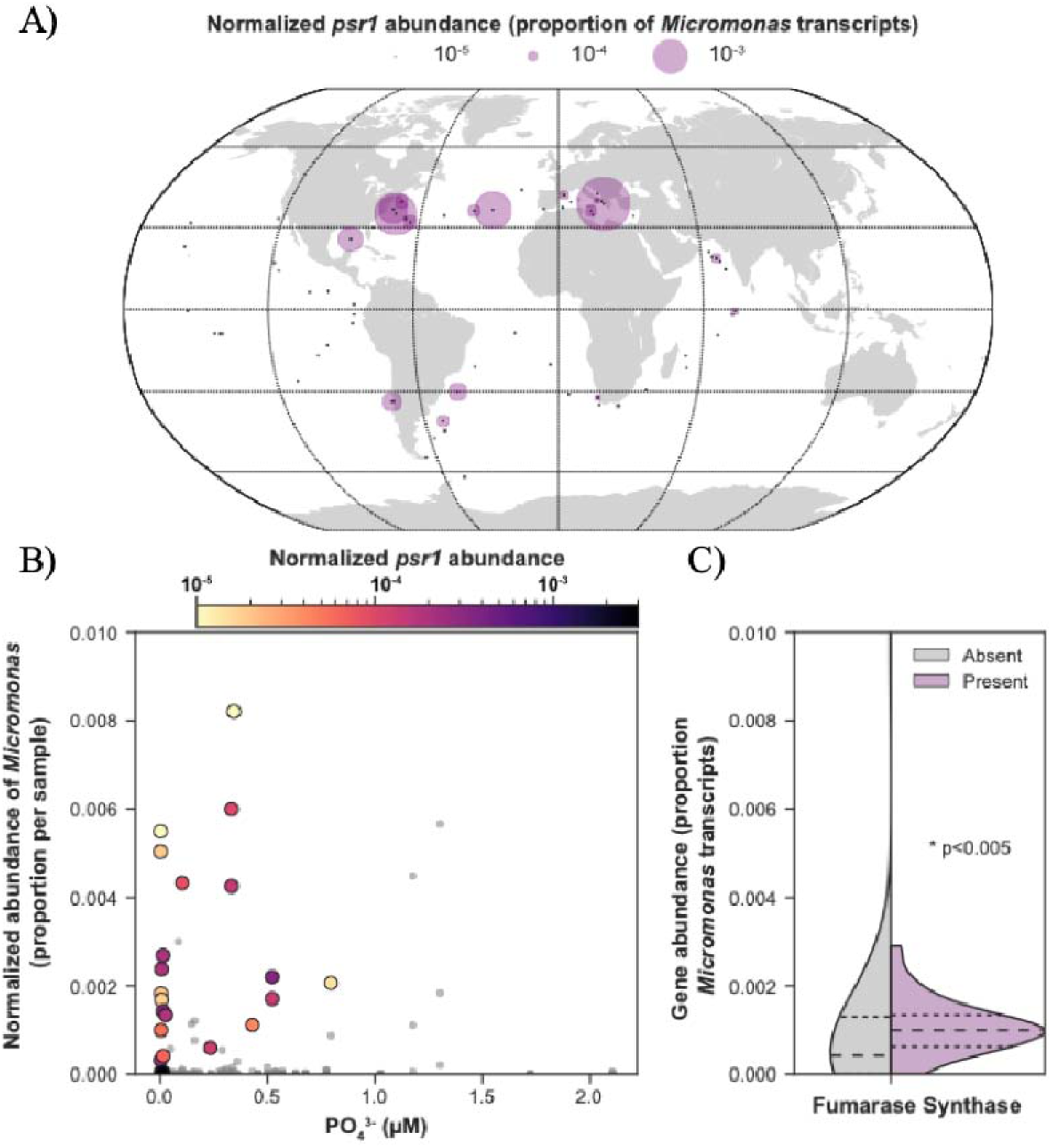
Geographic distribution of *Micromonas psr*1-like genes and relationship with phosphate concentration in the *Tara* Oceans eukaryotic gene atlas (Carradec, Pelletier, *et al*., 2017). The geographic distribution and relative abundance of *Micromonas psr*1-like genes in surface samples are depicted. Black dots represent stations where *Micromonas* was detected; the size of purple circles represents the relative abundance of *psr*1-like gene, where the FPKM of the psr1-like gene was normalized to the total FPKM of all *Micromonas*-associated transcripts (A). Relative abundance of *Micromonas* is plotted against phosphate concentrations for surface stations where *Micromonas* was detected. Grey circles indicate samples where *psr*1-like genes were not detected, while samples where psr1-like genes were detected are colored based on the relative expression of *Micromonas psr*1-like gene, again normalized to total *Micromonas* transcript abundance. No significant relationship was observed between the phosphate concentration and abundance of *Micromonas psr*1-like transcripts (*t*-test, *p* > 0.05) (B). A split violin plot depicting the distribution of expression of *Micromonas* fumarase synthase genes as a function of presence or absence of *psr*1-like gene. The larger dashed line represents the distribution mean and the smaller dashed lines represent the 25% and 75% quartiles. Significant difference in the distribution of expression between the presence and absence is assessed with Kolmogorov-Smirnov test (*p* < 0.005) (C).

Transcripts from MMETSP and *Tara* Oceans were generally short sequences and often contained one of the two characteristic domains for *psr*1-derived amino acid sequences (extended Results in the supplement). Field-derived transcripts of the *psr*1-like gene were geographically dispersed in the *Tara* Oceans dataset with the highest relative expression in surface samples and in the North Atlantic Ocean and Mediterranean Sea (Fig 5a). The majority of these transcripts from the *Tara* Oceans dataset occurred in samples with low phosphate concentrations (≤ 0.5 µM), but there was no significant relationship between *psr*1 expression and phosphate concentration (*t*-test, p < 0.05; Fig. 5b). We observed a significant positive relationship (Kolmogorov-Smirnov, p < 0.05) between the presence of *Micromonas psr*1-like genes and the relative expression of fumarase, a TCA cycle gene in *Micromonas*, which was up-regulated in cultures of *M. pusilla* (Bachy et al. 2018) and *M. commoda* (Whitney & Lomas 2016) under P-deficiency (Fig. 5c, see extended Discussion in the supplement). We did not find genes with significant homology to the *M. pusilla* putative *psr*1-like gene, in genomes or transcriptomes, within diatoms, the Heterokonta superphylum or the cryptophytes.

### 3.3 Putative regulatory element in Micromonas pusilla genes of interest

MEME analysis yielded a significant motif in *M. pusilla* genes involved in central carbon metabolism, lipid metabolism, and nucleotide metabolism (Fig. S5). The conserved motif is similar in nucleotide sequence to several TF-binding sites in *A. thaliana* (Table S4, Fig. S6; Franco-Zorrilla et al. 2014), including myb-like TF-binding motifs. The presence of a myb-like TF-binding motif is notable as the Psr1 protein contains a myb-like DNA binding domain (Wykoff et al. 1999). Several iterations of this analysis using genes not tied to the metabolites of interest yielded either no significant motif or motif sequences lacking similarity to myb-like domains. We observed a similar conserved motif in author-defined DE transcripts (Bachy et al. 2018) in *M. pusilla* CCMP1545 under P-deficiency (Table 1, Fig. S5). While the actual nucleotide sequence recognized by a TF must be experimentally determined (Franco-Zorrilla et al. 2014), the presence of a significant motif in each model and only in the expected genes, indicates that this conserved motif is likely biologically significant in *M. pusilla*. Specifically, this motif, with similarity to myb-like domains in *A*. *thaliana*, may represent a regulatory element where the Psr1-like protein binds.

### 3.4 Comparison of metabolomics-based predictions of P-responsive genes in *M. pusilla* CCMP1545 to transcriptomics analysis

We compared the author-defined DE transcripts in *M. pusilla* CCMP1545 to gene predictions based on our metabolomics data. As Bachy and colleagues (2018) used a more recent gene model than our original analysis, we confirmed the identity of the *psr*1-like gene in the updated model (CCMP1545/MBARI_models(ver_1), Table 2). We then analyzed a similar set of genes for a regulatory element where the Psr1-like TF might bind (Fig. S5), including genes that were DE in the P-deficient transcriptome relative to the replete (Table S1 from Bachy et al. 2018). Under P-deficiency, the *psr*1-like gene (JGI gene ID 360) was highly up-regulated (log-2 fold-change = 4.3, *q* = 0.00037) and nearly all of the predicted P-responsive genes were DE (Table 1). For example, *M. pusilla* up-regulated many genes in the TCA cycle but not pyruvate carboxylase (gene ID 6984). Genes involved in nucleotide and lipid metabolism (e.g., nucleoside phosphatase, long-chain acyl-coA synthetase) were DE and contained a conserved motif with similarity to myb-like regulatory element in *A. thaliana* (Table 1, Fig. S5). Lastly, the POX gene described in *E. huxleyi* (Rokitta et al. 2016) and observed in the archived gene model for *M. pusilla*, was absent, or not similarly annotated in the updated gene model. Instead, a copper amine oxidase (gene ID 5863) contained a significant conserved motif (Table 1, Fig. S5) and was DE under P-deficiency (Bachy et al. 2018).

We conducted a similar analysis with published transcriptome data from *M. commoda* (Whitney & Lomas 2016). The *psr*1-like gene (JGI gene ID 60184) in *M. commoda* was significantly up-regulated in the P-deficient treatment (defined by Whitney & Lomas 2016, Table S5), and we found a significant conserved motif in 18 DE genes. The significant motif identified in *M. commoda* differed in sequence from that discovered in *M. pusilla* and occurred in a different set of genes (see extended Results in the supplement).

## 4. Discussion

Our combined metabolomics and comparative genomic analyses of the response of *M. pusilla* CCMP1545 to P-deficiency has led to three conclusions: 1) there is a shift in intracellular metabolite composition, 2) a *psr*1-like gene is expressed in *M. pusilla* and other marine phytoplankton, and 3) the genes regulated by the Psr1-like protein may differ amongst algal species.

### 4.1 Micromonas pusilla exhibits a metabolic shift in response to P-deficiency

The general decrease in concentration of extracellular metabolites of P-deficient cells indicates reduced release of these central metabolites under P-deficiency. This is expected as nutrient-limited microalgae may have lower photosynthetic activity (Loebl et al. 2010, Halsey et al. 2014, Guo et al. 2018) and hence, produce less metabolic overflow. In contrast, most intracellular metabolites did not differ significantly in concentration between treatments. Because these metabolites represent many central metabolic pathways, it appears that *M. pusilla* does not respond to P-deficiency through a global decrease in metabolic activity, as was observed for the diatom, *Phaeodactylum tricornutum* (Feng et al. 2015). Interestingly, we found examples of functionally-related metabolites that behaved differently. For example, glutamate, proline, and methionine were higher on average, although variable, under P-deficiency, while phenylalanine and tryptophan were similar. In the TCA cycle, malate was variable but higher in concentration on average under P-deficiency, while citrate was lower on average and succinate was relatively unchanged. The opposing responses of these TCA intermediates was surprising given that others (e.g., Hockin et al. 2012, Goncalves et al. 2013) have observed a coordinated response to nutrient limitation in transcripts or proteins involved in the TCA cycle.

We observed additional non-uniform metabolite responses, with vitamins and glucosamine-6-phosphate higher on average under P-deficiency, but other P-containing compounds (e.g. glycerol-3-phosphate) and sugars were depleted. Previous work has suggested that organisms placed under nutrient stress are capable of diverting carbon flux between pathways (reviewed in Markou & Nerantzis 2013). Changes in carbon flux through the cell could result in coordinated abundance shifts in seemingly unrelated metabolites, as observed in this study. If the shunting of carbon is a coordinated response within *M. pusilla*, then we would expect genes involved in the synthesis and/or catabolism of these compounds to be co-regulated, and potentially mediated by a specific transcription factor. Indeed, in *M. pusilla,* we discovered a gene with significant homology to the transcription factor phosphorus starvation response gene (*psr*1) originally described in the freshwater alga *Chlamydomonas reinhardtii* (Wykoff et al., 1999).

### 4.2 Presence and expression of the transcription factor gene, psr1, in marine algae

In *C. reinhardtii* and in many plants, a *psr*1 gene encodes for a transcription factor (Psr1) that coordinates the metabolic response to P-deficiency in these organisms (Wykoff et al. 1999; Mosely et al. 2006). We detected a putative *psr*1-like gene in the *M. pusilla* genome and in the genomes and transcriptomes of major algal lineages (e.g., prasinophytes, dinoflagellates). The *psr*1-like genes that we discovered were generally not annotated as *psr*1 with the exception of *Ostreococcus tauri* (Derelle et al. 2006), although to our knowledge the potential role of this gene in the metabolism of *O. tauri* or other marine algae has not been discussed. The *M*. *pusilla psr*1-like gene contains two myb domains similar to those in *psr*1 from *C. reinhardtii* (Wykoff et al. 1999), and comparable to those in *phr*1 in *A. thaliana* (Rubio et al. 2001). No *psr*1-like gene was identified by our queries of diatom transcriptomes or genomes, suggesting that a phosphate starvation response gene akin to *psr*1 is present in diverse but phylogenetically constrained phytoplankton groups.

We found significant similarity between the *M. pusilla psr*1-like gene and transcript sequences from the MMETSP and *Tara* Oceans datasets, indicating that this gene is expressed by marine algae in cultures and in the environment. High-identity hits to sequences in the global ocean survey (GOS, Rusch et al. 2007) and *Tara* Oceans datasets underscore this gene’s prevalence in the oceans, particularly in regions characterized by chronically low phosphorus concentrations. Short sequences in some MMETSP and *Tara* Oceans sequences did not include enough of the C-terminal end to capture the myb coiled-coil domain. Thus, further investigation is required to confirm whether the identified transcripts are from *psr*1-like genes. Several GOS sequences, previously described as protein of unknown function, had homology to both characteristic Psr1 domains, suggesting that they could be re-annotated as *psr*1-like genes. The GOS sequences are likely derived from prasinophytes (i.e., *Micromonas* spp., *Ostreococcus* spp.) and haptophytes (e.g., *E. huxleyi*), highlighting the need to elucidate the role of *psr*1 in these organisms.

### 4.3 Potential role of Psr1 in the marine algal response to P-deficiency

Transcription factors typically recognize a specific motif in DNA where the protein binds to regulate each gene. If such a motif is observed, one can hypothesize that a gene containing the motif is regulated by the protein. We observed a conserved and enriched motif across genes involved in nucleotide biosynthesis, the TCA cycle and glycolysis, carbon fixation, fatty acid metabolism, and phosphate transport or salvage (Fig. 6). The conserved motifs discovered in both *M. pusilla* and *M. commoda* had unique sequences and were present in a slightly different set of genes (see extended Discussion in the supplement). The motif detected in *M. pusilla* is similar in nucleotide sequence to a myb DNA-binding domain in *A. thaliana.* This similarity is potentially significant, because the *psr*1/*phr*1 genes in *C. reinhardtii* and *A. thaliana* contain a myb DNA-binding domain. Thus, we hypothesize that the conserved motif detected in the *Micromonas* species represents binding sites for the Psr1-like transcription factors (TF).

**Fig. 6.**
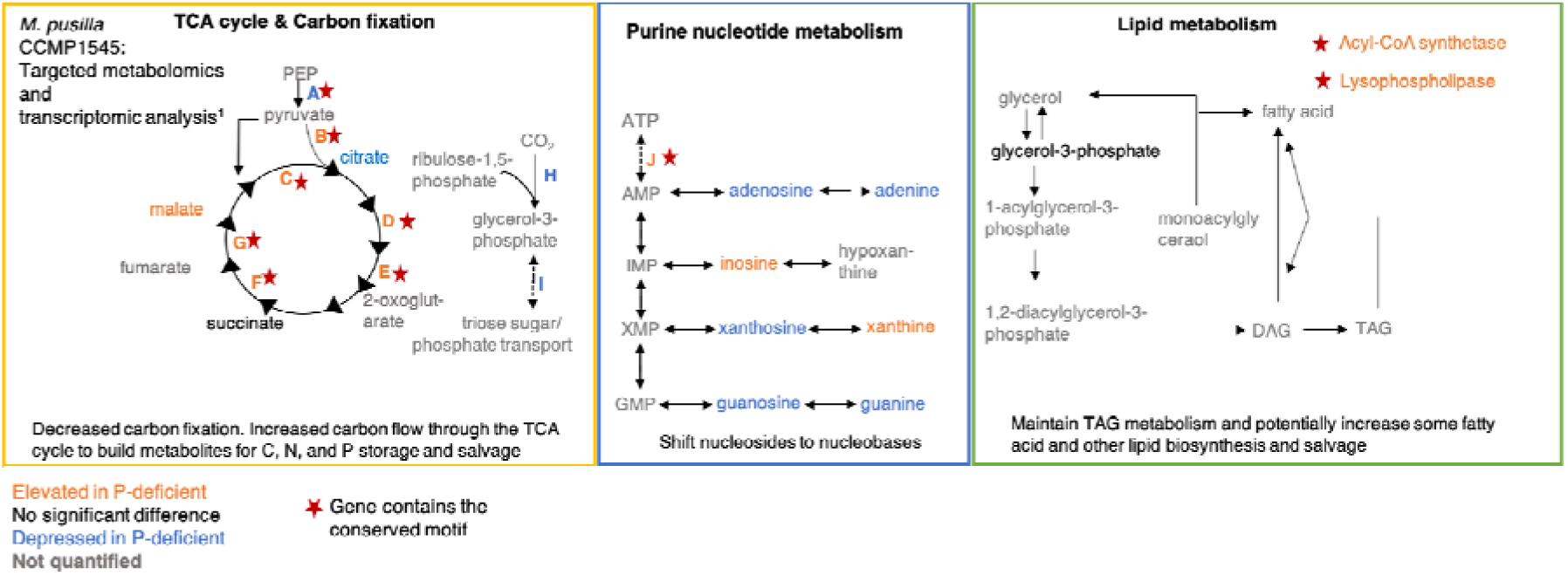
Conceptual model of Psr1-like TF regulation in *Micromonas pusilla* CCMP154 in three major metabolic pathways under phosphorus deficiency. The figure illustrates the relevant metabolites quantified in the present study and indicates if they are elevated (orange, ratio > 1 Figure 2), depressed (blue, ratio < 1 Figure 2) in concentration in the P-deficient cells relative to the P-replete cells. Some metabolite concentrations were equal in each treatment (black) or were not quantified (gray). Genes that contain the conserved motif with similarity to the myb-like regulatory element in *Arabidopsis thaliana* are marked with a red star. The orange and blue letters refer to elevated or depressed transcript levels for each gene (^1^data from Bachy *et al*., 2018). Descriptions of putative metabolic responses in each pathway are shown. Genes encode for: A) pyruvate kinase, B) pyruvate dehydrogenase, C) citrate synthase, D) aconitase, E) α-ketoglutarate dehydrogenase, F) succinate dehydrogenase, G) fumarase, H) ribulose-1,5-bisphosphate carboxylase, I) triose sugar transporter, J) nucleotide phosphatase DAG = diacylglycerol, TAG = triacylglycerol

The conserved motif was observed only in DE genes under P-deficiency by each *Micromonas* species (Bachy et al. 2018, Whitney & Lomas 2016), suggesting that these genes could be co-regulated. In bacteria, for example, genes within the well-characterized Pho-regulon all contain specific sequences (the PHO box) where the transcription factor binds to activate or repress the gene (Santos-Beneit 2015). These PHO box sequences are distinct among different bacterial species and strains. In *M. commoda*, the discovered motif was not significantly similar to myb-like DNA-binding motifs in *A. thaliana* and was not similar to the motif discovered in *M. pusilla*. Although we expected that the myb domain of the Psr1-like proteins in the two species would interact with similar binding regions in the genome, *psr*1-like derived amino acid sequences from *M. pusilla* and *M. commoda* were only 47% similar. TF protein sequences with up to 79% similarity have been shown to have distinct DNA-binding motif profiles (Franco-Zorrilla et al. 2014), so the two proteins in different species could reasonably have distinct DNA-binding motifs. If the Psr1-like TF regulates a different set of genes in the two *Micromonas* species, this may serve as a niche defining feature between these taxa. As further support for this hypothesis, the conserved motif was only observed in a subset of genes that were differentially expressed under P-limitation or deficiency (Whitney & Lomas 2016, Bachy et al. 2018). The observed differences in the *psr*1-like gene and differences in associated binding motifs may represent adaptation to distinct environments as *M. pusilla* was collected in the temperate English Channel, while *M. commoda* was isolated from the tropics (Worden 2006).

The *psr*1-like gene in *M. commoda* and *M. pusilla* was one of the highest DE genes in each experiment (Whitney & Lomas 2016, Bachy et al. 2018), suggesting that this potential TF plays a critical role in the P-deficiency response of *Micromonas*. Moreover, the *psr*1-like gene was highly expressed in two culture experiments with different growth conditions and sampled at different growth stages, suggesting a broad physiological relevance for the Psr1-like TF (Fig. 6). In contrast to the transcriptome results, a recent study describing proteome changes in *M. pusilla* CCMP1545 grown under P-limited conditions in a bioreactor (Guo et al. 2018), did not observe significant regulation of the Psr1-like protein (significance defined by Guo et al. 2018). The discordant results may be due to known differences in regulation between genes and proteins (Guo et al. 2018) or to different Psr1-like protein behavior under continuous culturing conditions. Regardless, these results warrant further investigation into the physiological role of the Psr1-like protein in *M. pusilla*.

Variable and non-uniform dynamics of several TCA cycle metabolites between treatments suggested that this pathway may be involved in the metabolic response to P-deficiency in *M. pusilla*. Indeed, nearly all of the genes involved in the end of glycolysis and in the TCA cycle were up-regulated in *M. pusilla* (Bachy et al. 2018) and these genes contained the conserved motif that may function as a regulatory element where the Psr1-like TF could bind. Fumarase, in particular, was upregulated in both *M. pusilla* and *M. commoda* transcriptomes under P-limitation or deficiency. If fumarase expression is regulated by the Psr1-like protein, expression of these two genes (fumarase and *psr*1) should be correlated in field populations. We observed support for this hypothesis within the *Tara* Oceans dataset, although we observe weaker relationships between Psr1-like gene expression and other up-regulated genes in either *M. pusilla* or *M. commoda* (see extended Discusssion in the supplement). In *M. pusilla*, carbon flow through the TCA cycle may increase as a means to fuel TAG or starch production and to pull potentially damaging energy away from the photosystems (Norici et al. 2011, Klok et al. 2013). Similarly, several proteins involved in glycolysis (e.g., pyruvate dehydrogenase) and the TCA cycle (e.g., succinate dehydrogenase) were found to be up-regulated in another study with *M. pusilla* under P-limitation (Guo et al. 2018), lending some support to this hypothesis. Insight from the latter study underscores potential differences in regulation of genes and enzymes involved in glycolysis and the TCA cycle (Guo et al. 2018). For example, we found congruency in up- or down-regulation of some proteins from Guo et al. (2018) that we would expect in *M. pusilla* under P-deficiency based on our combined comparative genomics and metabolomics approach (i.e., pyruvate kinase, pyruvate dehydrogenase, oxoglutarate dehydrogenase, and succinate dehydrogenase) but not others (i.e., fumarase, aconitase, citrate synthase). Analysis of transcript, protein, and metabolite responses in *M. pusilla* under the same growth conditions will be required to elucidate the role of Psr1 in this and other taxa.

In addition to TCA cycle intermediates, we observed shifts for several purine nucleosides between treatments and detected the conserved motif in genes for a nucleoside phosphatase and aspartate transcarbamylase. These results suggest increased nucleotide salvage through purine nucleotides and nucleosides, a mechanism described in *M. commoda* (Whitney & Lomas 2016) and in other phytoplankton (Dyhrman & Palenik 2003, Kujawinski et al. 2017). Previous phytoplankton work has also highlighted expression of proline oxidase (POX) (Rokitta et al. 2014, Rokitta et al. 2015) in nutrient limitation experiments (including P-deficiency) with the haptophyte *E. huxleyi,* where this enzyme may have a role in stabilizing the mitochondrial membrane and in detecting cellular nitrogen levels. The POX gene was not present in *M. pusilla* but was up-regulated in *M. commoda* (Whitney & Lomas 2016, Table S4). In *M. pusilla*, a copper amine oxidase was up-regulated (Bachy et al. 2018) and contained the conserved motif that may function as a regulatory element for the Psr1-like TF. Thus, POX and copper amine oxidase represent gene targets for further investigation.

Maat et al. (2014) noted that both *E. huxleyi* (Borchard et al. 2011) and *M. pusilla* (Maat et al. 2014) exhibited a similar metabolic response to elevated *p*CO_2_ levels and P-deficient conditions. While there may be multiple factors underlying the response of these organisms, both contain the *psr*1-like gene which could direct part of a shared metabolic response to elevated *p*CO_2_ and nutrient-limited conditions. While transcriptomic analysis of *E. huxleyi* was not performed in the study by Borchard et al. (2011), Rokitta and colleagues (2014, 2016) performed microarray transcriptome analysis of *E. huxleyi* cells under separate nitrogen- and P-deficient conditions (normal pCO_2_). The transcripts were only partial sequences; however, two up-regulated genes detected by Rokitta et al. (2016), contained myb-like DNA binding domains (gene IDs: GJ02872, GJ03978). Targeted experimental work is necessary to elucidate the presence and role of a *psr*1-like gene in *E. huxleyi*. If there is a common role for the *psr*1-like gene in *E. huxleyi* and *M. pusilla* in their response to nutrient limitation and elevated pCO_2_, this could have important implications for predicting the physiological response of the wide range of organisms that contain this *psr*1-like gene to changing environmental conditions.

The potential impact of Psr1-regulated genes on the ecological roles of *Micromonas* spp. is unknown, but our results suggest that the Psr1-like protein coordinates a metabolic shift in these organisms under P-deficiency, altering the intracellular flow of carbon and other elements. More comprehensive examination of these metabolic responses, which likely vary to some extent among organisms, will be paramount to improving models of trophic carbon flow. More experiments are needed to characterize the structure and role of the *psr*1-like gene in *Micromonas* spp. and other phytoplankton, including: (1) confirming the presence of the *psr*1-like gene with genetic experiments, (2) determining the conditions under which the Psr1-like protein is abundant, (3) identifying the taxon-specific genes affected by the Psr1-like protein, (4) verifying the interaction of the Psr1-like protein with the hypothesized binding sites, and (5) comparing the genetic and metabolic responses to P-deficiency between organisms containing the *psr*1-like and those that do not. Exploring the underlying biology of the *psr*1 gene will facilitate mechanistic understanding of the complex response of these organisms to P-limitation and will enhance our ecological and biogeochemical predictions.

## Acknowledgements

The authors would like to thank Krista Longnecker for performing the TOC analysis and Matthew Johnson and Elizabeth Harvey for assistance with flow cytometry and FIRe analysis. This research was funded by the Gordon and Betty Moore Foundation through Grant GBMF3304 to E. Kujawinski.

