## Supplemental File 1 for "A phosphate starvation response gene (*psr*1-like) is present and expressed in *Micromonas pusilla* and other marine algae"

**Extended Methods:**

*Assessment of culture contamination*

DAPI-staining was performed using 60 μl of culture fixed with 60 μl buffered 37% formaldehyde (buffered with sodium borate) and frozen at -80°C. The fixed cells were stained with 4’,6-diamidino-2-phenylindole dihydrochloride (DAPI, 70 μg ml^-1^ final concentration) on Isopore membrane filters (0.2 μm, Millepore). We examined the filters using a Zeiss Axiostar Plus microscope, where 25-35 fields or at least 150 algal cells were counted as described in Fiore *et al*. (2015). No contamination was found at any time point.

*TOC analysis*

Samples for TOC analysis were brought up to 40 ml volume with MilliQ water, acidified to pH = 3 and stored at 4°C until analysis with a Shimadzu TOC-V_CSH_ total organic carbon analyzer (Hansell & Carlson, 2001). Standards provided by Prof. D. Hansell (University of Miami) were used for instrument calibration and were made fresh each day as described in Longnecker (2015). Nutrient analysis was performed at the Nutrient Analytical Facility at WHOI using a SEAL AA3 four-channel segmented flow analyzer according to USEPA approved protocols to determine concentrations of ammonium (NH_4_^+^), nitrate plus nitrite (NO_x_^-^), and phosphate (PO_4_^-3^).

*Metabolite extraction and instrument method*

We processed the sample filtrate using solid phase extraction (SPE) with 1 g/ 6 cc PPL cartridges (Agilent, Santa Clara, CA) to concentrate extracellular metabolites and to remove salt (Dittmar *et al*., 2008). Metabolites were eluted from the column using 100% methanol and these extracellular samples were stored at -20°C until analysis. Just prior to analysis on the instrument described below, the methanol extract was dried down and re-constituted in 500 μl of 95:5 water:acetonitrile and deuterated biotin (d2-biotin) was added to each sample as an HPLC injection standard (final concentration 0.05 μg ml^-1^). At this stage, we used 100 μl of the extracellular extract for targeted metabolomics analysis.

A pooled sample for quality control consisted of 10 parts intracellular extract and one-part extracellular extract of each experimental sample. The contributions of intracellular and extracellular samples to the pooled sample (i.e., 10:1 described above) were determined by analyzing several ratios of intracellular to extracellular samples using liquid chromatography coupled with Fourier transform ion cyclotron resonance (FT-ICR) mass spectrometry as described by Kido Soule *et al*. (2015). Non-metric multidimensional scaling (nMDS) was used to determine the ratio at which the profile of the pooled sample is similar to both the extracellular and intracellular metabolic profiles, indicating that the pooled sample is representative of both intracellular and extracellular samples. These pooled samples were run at the beginning of the analytical run and approximately every six samples and then at the end, which we used a measure of consistency across the analytical time-frame. We did not see significant changes in metabolite peak intensity across the pooled samples, indicating that there were no ionization problems within the analytical time-frame.

*Genome comparisons and* Tara *Oceans data analysis*

The IMG genomes included the cryptophyte *Guillardia theta*, and several diatom species and other heterokonts: *Thalassiosira pseudonana*, *Phaeodatylum tricornutum*, *Fragilariopsis cylindrus, Pseudo-nitzschia multiseries*, *Aplanochytrium* *kerguelense*, *Aurantiochytrium limacinum*, *Aureococcus anophagefferens*, *Ochromonadaceae* sp., and *Pelagophyceae* sp. For the *Tara Oceans* dataset, metatranscriptome assemblies and corresponding metadata were downloaded from the Tara Oceans Eukaryotic Gene Catalog (Carradec, Pelletier *et al*., 2017; http://www.genoscope.cns.fr/tara/). The unigene sequence files were used to create a blast database and were queried using with the *M. pusilla* CCMP 1545 *psr*1-like-derived amino acid sequence (JGI ID: 61323). BLAST results were compiled with sample metadata, parsed with Dask (Dask Development Team), and visualized using the Matplotlib Basemap Toolkit. As the published metatranscriptomic dataset was in fragments per kilobase million (FPKM), expression values were not directly comparable across samples. Therefore, we selected only the *psr*1-like unigenes that were assigned that taxonomic classification of “*Micromonas*” and normalized the expression value of the *psr*1 unigenes to the total abundance of all unigenes that were taxonomically classified as *Micromonas* per sample. The Jupiter notebook workflow for this work is available at https://github.com/AlexanderLabWHOI/micromonas-psr1-tara. Corresponding phosphate data were obtained from Carradec, Pelletier, *et al*. (2017) supplemental table 5.

*Comparison to* Micromonas commoda *RCC299 gene expression under P-deficiency*

We performed the same protocol for regulatory element motif discovery and identification on gene sequences from *M. commoda* RCC299 (gene model: *Micromonas pusilla* NOUM17/FrozenGeneCatalog_20090404) and CCMP1545 (gene model: *Micromonas pusilla* CCMP1545/MBARI_models(ver_1)). We first queried the DE genes in *M. commoda* RCC299 exposed to P-deficient conditions (Whitney & Lomas, 2016) for the *psr*1-like gene as well as for genes that we expected to be regulated by the Psr1-like TF based on our *M. pusilla* observations and the literature (e.g., Sharma *et al*., 2012; Rokitta *et al*., 2016).

**Extended Results:**

*Growth of* Micromonas pusilla *CCMP1545* *under phosphorus deficiency*

Inorganic nutrient concentrations decreased over time in the media with the exception of a small increase in ammonium in both treatments during stationary phase (Figs. S7a,b,c,d). Photochemical efficiency decreased in late exponential growth in both treatments although it was maintained around 0.4 in the P-replete cultures in stationary phase (Fig. S8a). Cell size, as estimated by forward scatter, increased during the experiment for P-deficient cells only (Fig. S8b).

*Search for* psr*1 in the GOS, MMETSP, and* Tara *Oceans datasets*

The putative *psr*1-like transcripts from the MMETSP dataset all contained the myb DNA-binding domain, but only 55 of the 148 matches also contained the myb coiled-coil domain at the C-terminal end of the protein sequence (Table S2), indicating that further investigation into the identified genes is required to determine if they are psr1-like genes or potentially other transcription factors. The sequences in the GOS dataset identified as significant matches to *psr*1 were most similar to *Ostreococcus* spp., *Chlorella* *variabilis*, and the haptophyte *Pleurochrysis carterae* (Fig. 4). Of the 252 hits in the *Tara* Oceans assemblies, 199 contained the myb-like DNA binding domain, while only 21 contained this DNA binding domain and the myb coiled-coil domain at the C-terminal end of the protein sequence, similar to the MMETSP results. Transcripts of the *psr*1-like gene derived from prasinophytes were detected in the Pacific, Atlantic, Indian, and Southern Oceans and in the Mediterranean Sea (Fig. 6a). The *Tara* Oceans project analyzed metagenome and metatranscriptome assemblages for different filter fractions and at three depths (Sungawa *et al*., 2012; Carradec, Pelletier *et al*., 2017) and we detected the highest number of hits to the *psr*1-like gene within the 0.8 – 5 µm filter fraction (data not shown) and at the surface (Figs. 6a, b). Lastly, we observed transcripts from the *Tara* Oceans dataset generally at or below 2 µM phosphate in the field (Fig. 6b). One of the identified genes (from *O. tauri*) was already annotated as *psr*1 (Derelle et al., 2006), while candidate *psr1-like* genes in other organisms were annotated as hypothetical protein or with a myb-like DNA-binding domain but not as *psr*1.

*A* psr*1-like gene and the search for significant motif in* Micromonas commoda *RCC299*

In *M. commoda*, the *psr*1-like gene (JGI gene ID: 60184) was significantly up-regulated in the P-deficient treatment (Table S4), and we found a significant conserved motif in 18 DE genes in *M. commoda*. The significant motif identified in *M. commoda* differed in sequence from that discovered in *M. pusilla* and occurred in a different set of genes*.* However, putative binding motifs for Psr1 are present in the same pathways in the two *Micromonas* species and are similar to other DNA-binding motifs in *A. thaliana* (Fig. S6, Results S2). The *psr*1-like genes in RCC299 and CCMP1545 exhibited relatively low homology to each other but were equally similar to the *psr*1-derived amino acid sequence in *C. reinhardtii* (Table S5).

Our query of the *M. commoda* P-deficient transcriptome (defined by Whitney & Lomas, 2016) indicated that several, but not all of the genes that we expected to be DE, based on our metabolomics analysis in CCMP1545, were significantly expressed in the P-deficient treatment (Table S4). For example, in the TCA cycle, fumarase (96474), the enzyme that produces malate from fumarate, was upregulated, while citrate synthase (96682), which produces citrate from acetyl-coA and oxaloacetate, was downregulated. Additionally, malate dehydrogenase (75917), which converts malate to oxaloacetate (i.e., removes malate) was also upregulated in *M. commoda*. Our metabolomics analysis with *M. pusilla* indicated non-significant differences in the concentration of malate and citrate between treatments (Fig. 2), which may be related to the expression of the corresponding biosynthetic genes. However, we are cautious in our interpretation here as these metabolites are involved in many reactions that could influence their intracellular concentrations and the gene expression may differ between the two *Micromonas* species. Several genes involved in fatty acid and glycerolipid (including TAG) biosynthesis were also up-regulated in *M. commoda* under P-deficient conditions, including a phosphatidic acid phosphatase (58855), which catalyzes the second to last step in TAG production, and acylglycerol lipase (80225), involved in lipid catabolism (e.g., membrane lipids) (Table S3). Similarly, the POX gene (100385) was also significantly upregulated in the P-deficient treatment in *M. commoda*. Unexpectedly, the gene encoding for aspartate carbamoyltransferase (98407), which catalyzes the first step in pyrimidine biosynthesis using aspartate as a substrate was not downregulated in RCC299. We hypothesized that higher average concentrations of aspartic acid in P-deficient *M. pusilla* might be a result of down-regulation of *de novo* synthesis of pyrimidines, the first step of which uses aspartic acid.

We hypothesized that if the Psr1-like TF is functioning in *M. commoda* under P-deficient conditions, then a conserved motif similar to that identified in *M. pusilla* with similarity to myb-like DNA-binding domains would be present in Psr1-regulated genes in *M. commoda*. We first used a set of genes in *M. commoda* that were homologous to those analyzed in *M. pusilla* to search for a conserved motif. This search yielded no significant motifs. Subsequently, we used DE genes from *M. commoda* (those that were DE and in the pathways we highlighted from *M. pusilla*) for motif discovery, yielding a significant motif different from that discovered in CCMP1545 but similar to other DNA-binding motifs characterized in *A. thaliana* (Fig. S6; Franco-Zorilla *et al*., 2014). Our metabolomics analysis and results of the motif discovery in each strain combined with the published transcriptomic data, indicate that the genes regulated by the Psr1-like TF may vary between *M. pusilla* and *M. commoda* (Fig. 6).

**Extended Discussion:**

*The potential role of* psr*1 in Micromonas*

In comparing the transcriptome responses of *M. pusilla* and *M. commoda* to limitation of P, it is difficult to assess whether observed differences in gene expression are a result of divergent metabolic responses to P-limitation or from differences in experimental design between the two experiments. Yet, the comparison we discuss here provides multiple testable hypotheses for future work. Based on previous studies, we expected genes involved in triacylglycerol (TAG) production to increase under P-deficiency but found divergent responses between the two species. Two TAG production genes were up-regulated in *M. commoda*, including the penultimate step in TAG production (putative phosphatidic acid phosphatase), which contained a putative regulatory element where the Psr1-like TF could bind. By contrast, neither of the two TAG metabolism genes that were DE in *M. commoda* were DE in *M. pusilla*; instead two different genes involved in fatty acid biosynthesis and metabolism were DE in *M. pusilla*. Additionally, a starch-binding protein was highly up-regulated (defined by Bachy *et al*., 2018) in the *M. pusilla* transcriptome under P-deficiency (gene ID 9633). It is possible that *M. pusilla* may invest resources in starch storage, similar to the model alga, *Chlamydomonas reinhardtii* (Baj), at least under the growth conditions used by Bachy *et al*. (2018).

As described in the main text, we also observed shifts in concentration for several purine nucleosides between treatments for *M. pusilla* and detected the conserved motif in the genes for a nucleoside phosphatase and aspartate transcarbamylase. Interestingly, while a 5’-nucleotidase was observed to be DE in *M. commoda* (Whitney & Lomas, 2106) under P-deficiency, it was not DE in *M. pusilla* (Bachy *et al*., 2018). In the *M*. *pusilla* transcriptome (Bachy *et al*., 2018), a nucleotide phosphatase was up-regulated and may function in a similar capacity in nucleotide salvage.

In contrast to *M. pusilla* (Bachy *et al*., 2018), only a few genes involved in the TCA cycle were up-regulated in *M*. *commoda*, including genes involved in malate and oxaloacetate (OAA) metabolism and a gene for pyruvate carboxylase (PC), which converts pyruvate to oxaloacetate. The expression of these TCA-cycle genes may reflect an increase in the malate-aspartate shuttle, a redox reaction that drives the production of NAD^+^ from NADH in the mitochondrial membrane. In *M. commoda*, it is possible that PC-based conversion of pyruvate to OAA combines with an increase in the malate-aspartate shuttle function as a stress response that allows continued glycolysis and ATP production *via* oxidative phosphorylation (Yoshida *et al*., 2007; Easlon *et al*., 2008). In this scenario, carbon flow into the TCA cycle would be limited and the genes controlling malate, aspartate, oxaloacetate, and pyruvate would be co-regulated. This hypothesis is supported by the observation of a conserved motif that may function as a regulatory element in the relevant genes and by preliminary gene expression analysis of one of the TCA cycle genes that was up-regulated in both *M. pusilla* and *M. commoda* transcriptomes under P-deficiency, the fumarase gene. If fumarase expression is regulated by the Psr1-like protein, expression of these two genes (fumarase and *psr*1) should be correlated in field populations. We observed support for this hypothesis within the *Tara* Oceans dataset (Fig. S9), where the expression of *psr*1 was positively and significantly correlated with the expression of the fumarase gene in *Micromonas*. However, we observed weaker relationships between *psr*1-like gene expression and other genes that were up-regulated in either *M. pusilla* or *M. commoda* only.

The other two genes that we tested in a similar manner to fumarase, are citrate synthase and pyruvate kinase, both are involved in the TCA cycle and were differentially regulated in *Micromonas* species under P-deficiency. Citrate synthase was up-regulated in *M. pusilla* and down-regulated in *M. commoda*, while pyruvate kinase exhibited the opposite behavior, it was down-regulated in *M. pusilla* and up-regulated in *M. commoda*. In contrast, fumarase was up-regulated in both species. In the *Tara* metatranscriptome dataset, citrate synthase and fumarase exhibited a significantly different distribution in samples with *Micromonas* *psr*1 gene expression than in samples without this expression, while pyruvate kinase was not significantly different (Komogorov-Smirnov, *p* < 0.005, Fig. S9). Citrate synthase and fumarase exhibited a higher average expression in samples with *psr*1 expression, corroborating our hypothesis that the *psr*1 gene product regulates these genes. We note that for all three of these genes there are multiple factors that regulate their expression; however, the observation of a significant relationship with *psr*1 expression provides intriguing evidence for the role of a Psr1-like transcription factor in central metabolism and the metabolic response of *Micromonas* to P-deficiency.

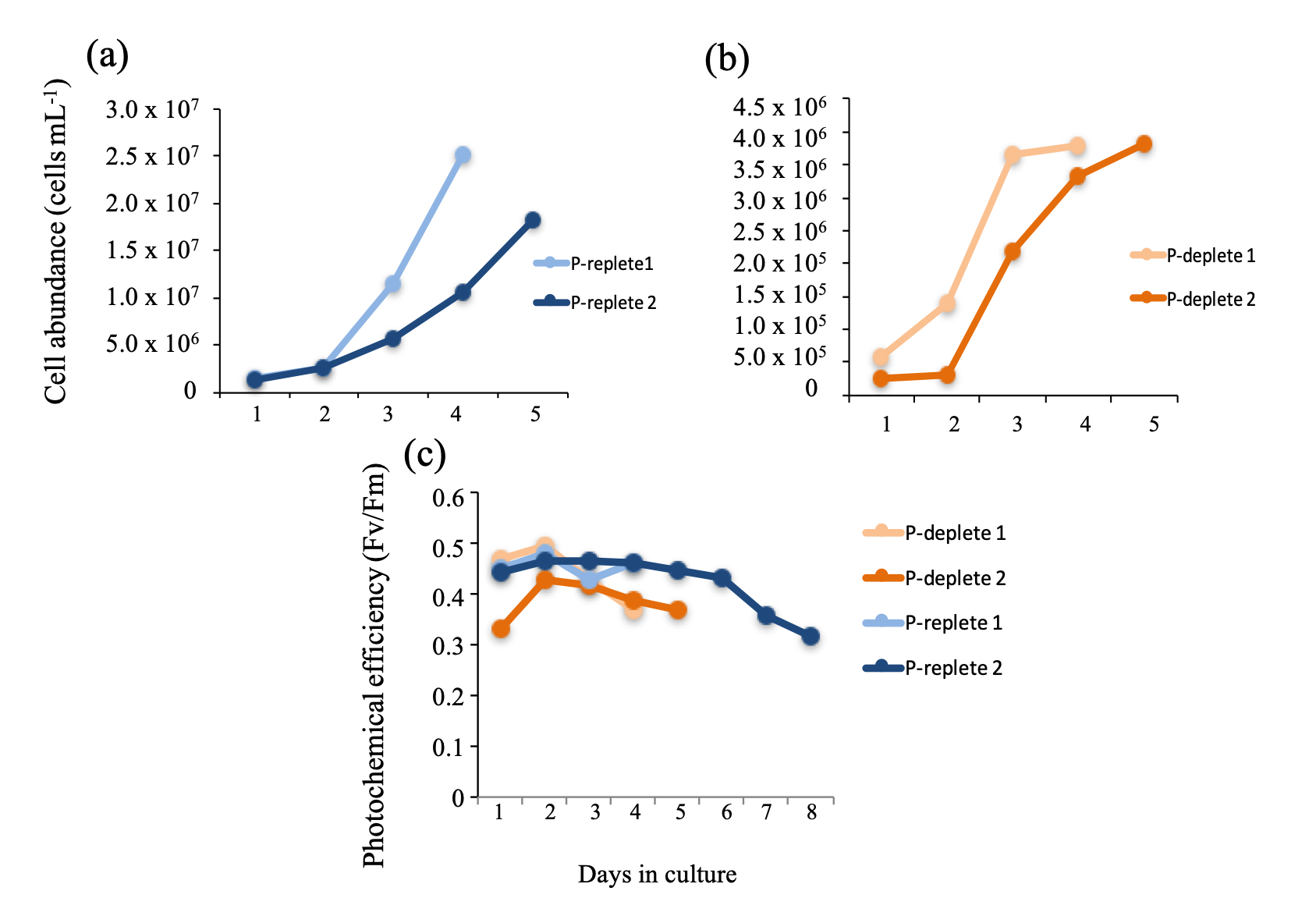

Figure S1. Growth information on *Micromonas pusilla* CCMP1545 cultures prior to experimental setup. Cell abundance over time for P-replete (blue, a) and P-deficient (orange, b) cultures and photochemical efficiency for both treatments (c). P-replete 1 and P-deficient 1 were used to inoculate P-replete 2 and P-deficient 2, respectively, and these in turn, were used to start the experiment.

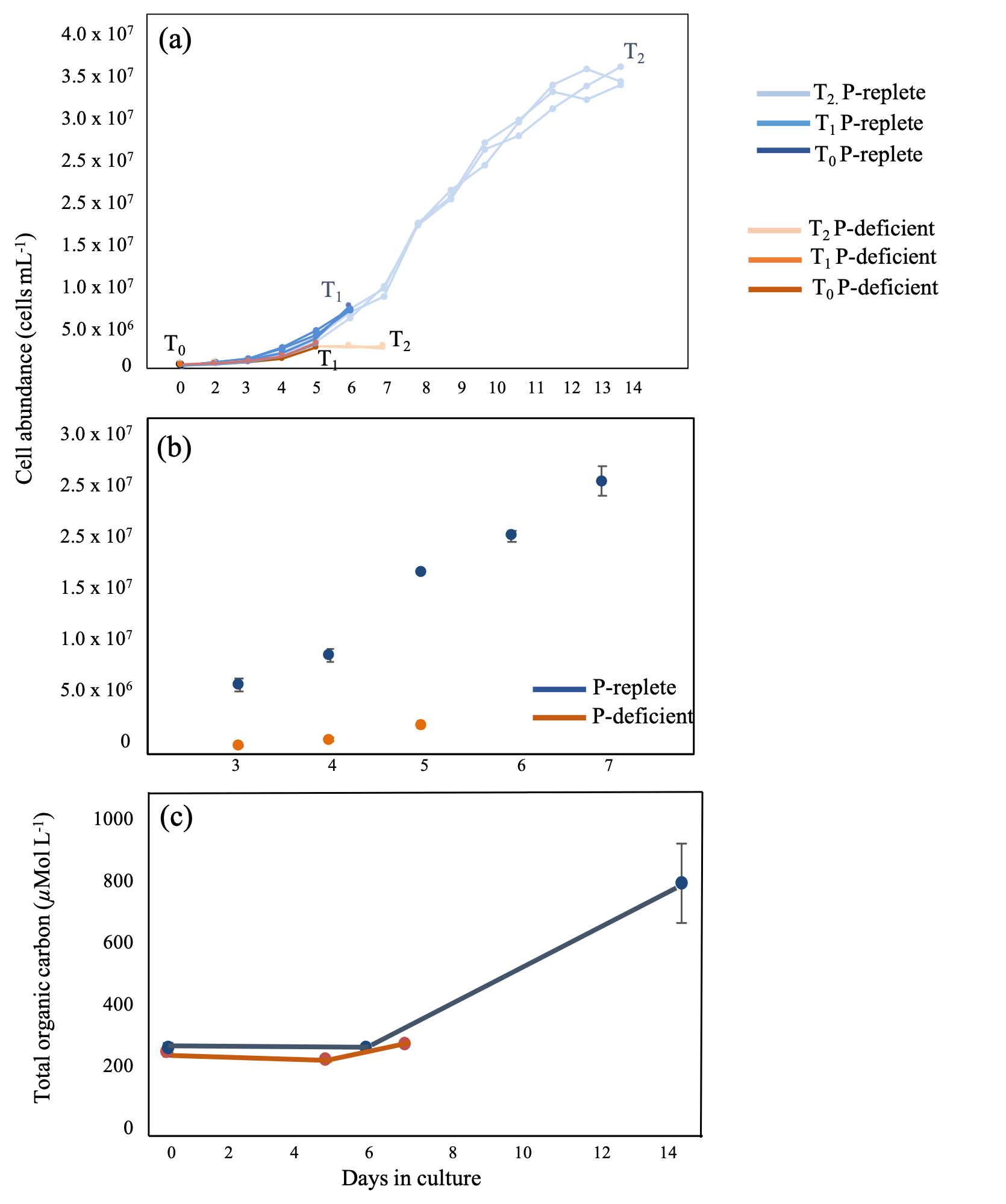

Figure S2. Growth curves for *Micromonas pusilla* CCMP1545 in P-replete and P-deficient media. Points represent the average of three replicates with standard deviation. Sampling points are indicated by the last point of each color and annotated as T_0_ (start of the experiment), T_1,_ and T_2_ in (a). The difference in scale between treatments during rapid growth is shown in (b). Cell abundance was quantified with flow cytometry (a, b); while growth could be monitored by total organic carbon accumulation (c).

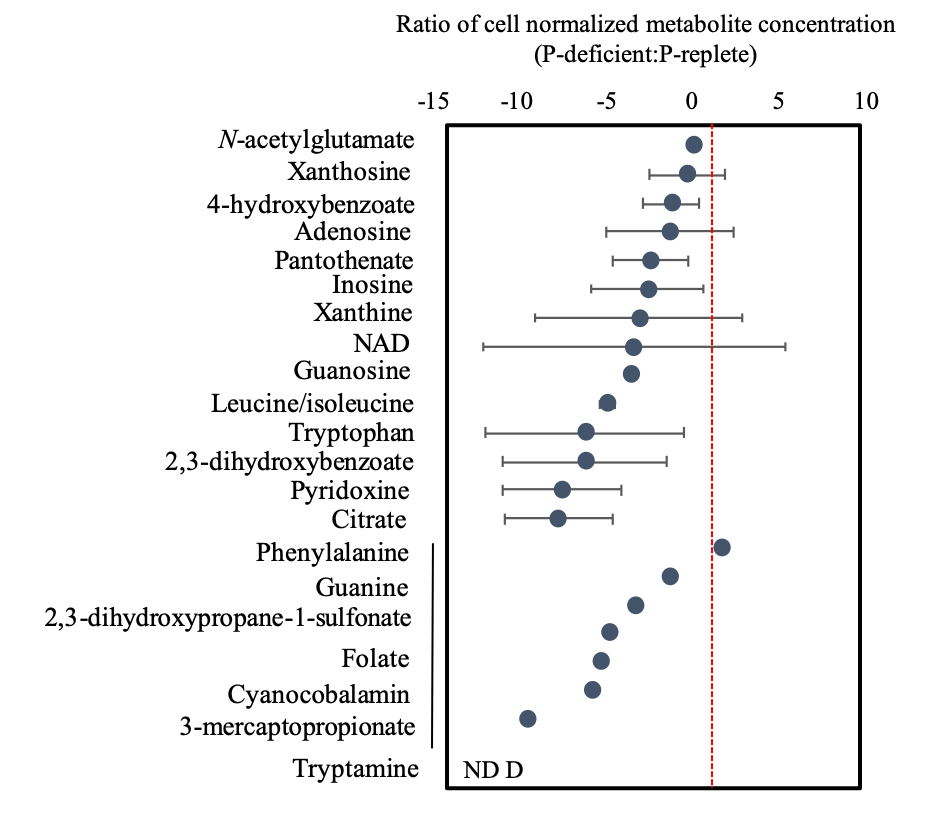

Figure S3. Average ratio of P-deficient to P-replete extracellular metabolite concentrations (log_2_) during stationary growth (T_2_). The extracellular metabolite profile in stationary phase represents a snapshot of the cumulative changes over time in the media. Concentrations are standardized to cell number in each treatment and standard deviations are shown. Metabolites are listed in order of descending concentration within three groups; those that were detected in enough replicates for standard deviation to be calculated, those where only one ratio could be calculated (black line), and those that were not detected (ND) in any of the three replicates for one treatment (D = P-deficient).

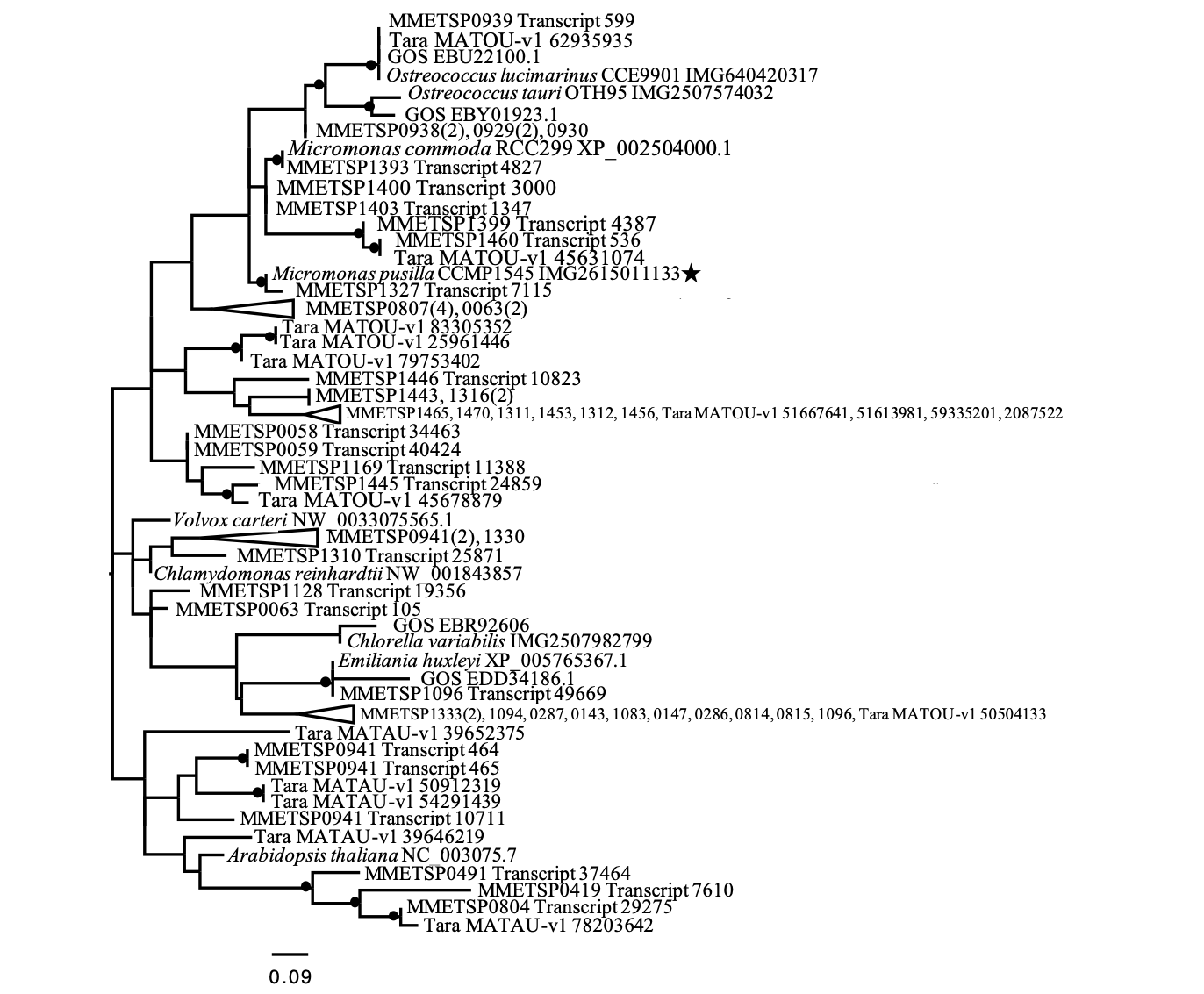

Figure S4. Occurrence and phylogenetic relationship of *psr*1-like genes in marine phytoplankton. All sequences that included the conserved SHLQKYR domain were included in the alignment (see Materials and Methods in main text and extended Methods for details). Maximum likelihood method based on the JTT matrix-based model (Jones *et al*., 1992). The tree with the highest log likelihood (-977.06) is shown and is based on the derived amino acid sequences for *psr*1 and *psr*1-like genes from eukaryotic phytoplankton. Bootstrap support for branches that are 50% or great are marked with a black circle. The tree is unrooted and drawn to scale, with branch lengths measured in the number of substitutions per site. The analysis involved 85 amino acid sequences. All positions containing gaps and missing data were eliminated. There were a total of 24 positions in the final dataset. Evolutionary analyses were conducted in MEGA7 (Kumar *et al*., 2016). Accession or IMG gene identification numbers are shown with the taxon name. For clarity, collapsed branches include MMETSP identification numbers but not transcript ID and number of sequences are shown in parentheses. *Micromonas pusilla* CCMP1545 used in this study is marked with a star.

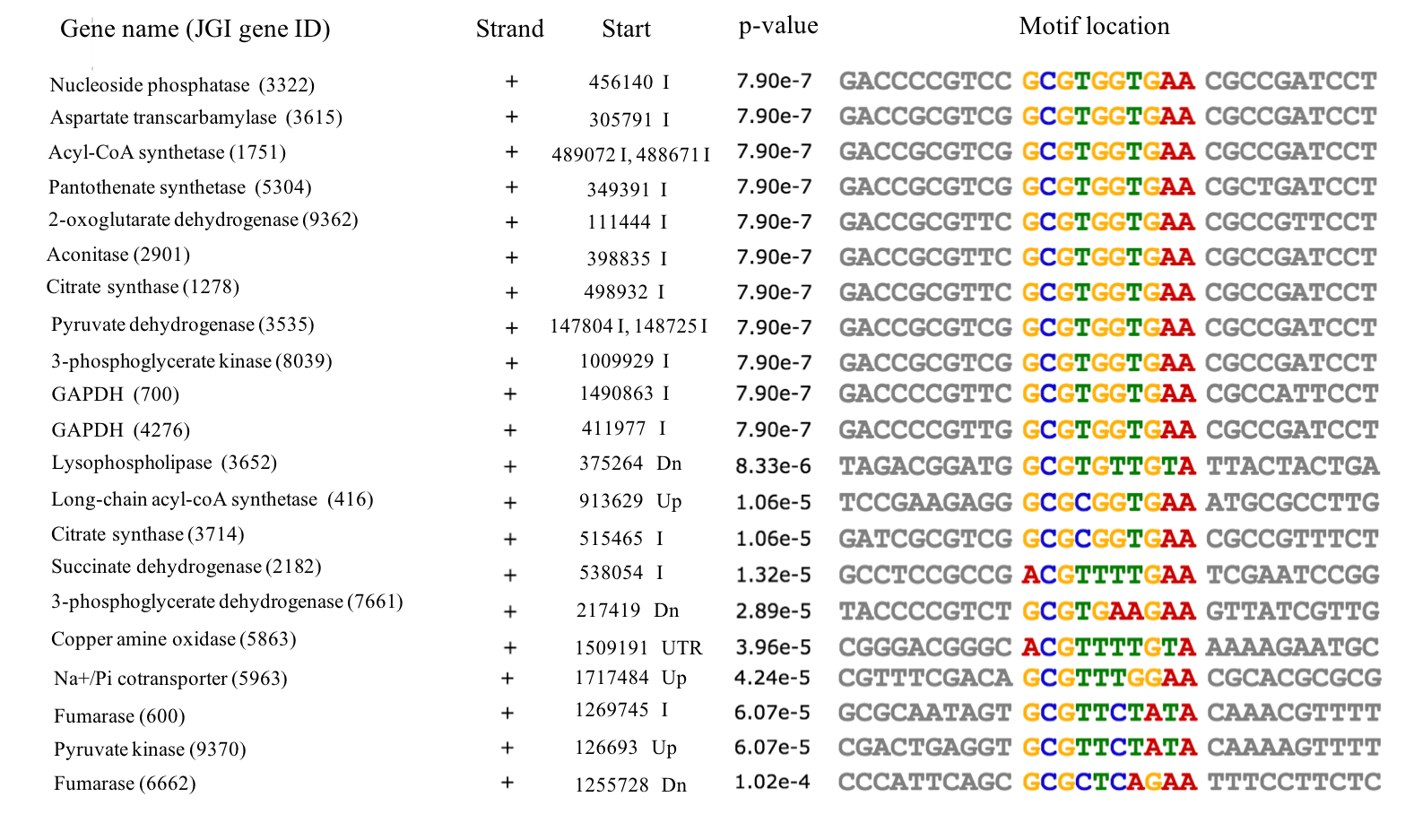

Figure S5. Overview of the significant motif in each gene in *Micromonas pusilla* CCMP1545. Genes are in order of increasing p-value. Each gene sequence is comprised of 500 nucleotides upstream (Up) of the gene, untranscribed regions (UTR), introns if present (I), and 500 nucleotides downstream (Dn) of the gene concatenated together. Shown here are the genes of interest with the JGI protein identification (MBARI_models (ver1)) number, the strand (+ or -) with the motif, start location of the motif in the gene, the p-value representing the probability of finding this motif by chance within the nucleotide sequence for that gene, and the location of the motif with in the concatenated nucleotide sequence.

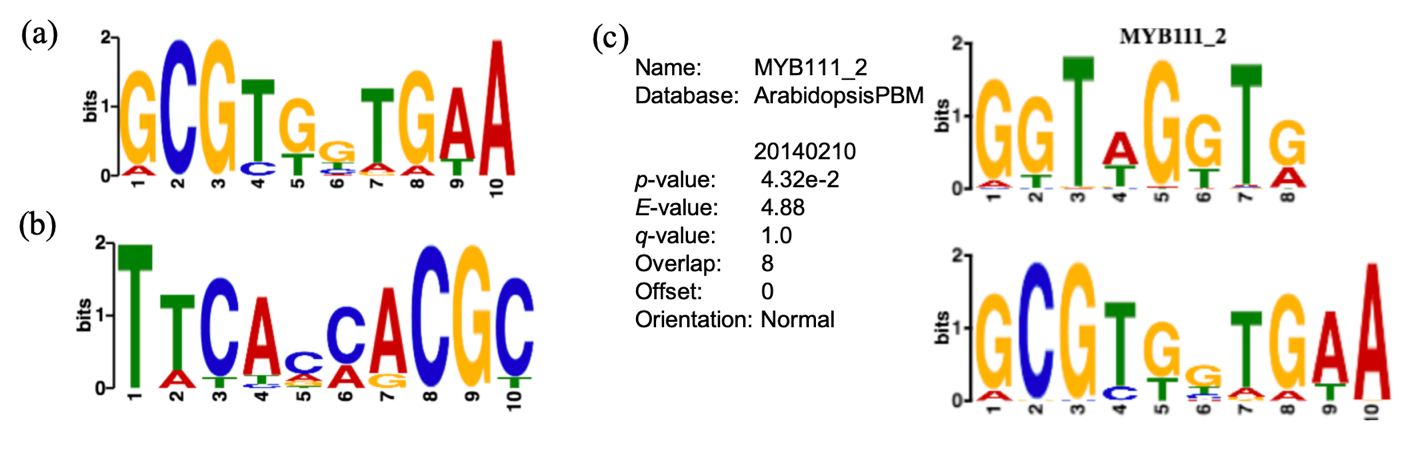

Figure S6. Significant motif present in *Micromonas pusilla* CCMP1545 gene sequences described in Table 1. The motif consensus logo for the positive, or coding strand (a), and for the negative strand (b) are shown. The consensus motif is statistically similar to the regulatory element MYB111_2 within the *Arabidopsis* regulatory element database (Franco-Zorilla *et al*., 2014) (c).

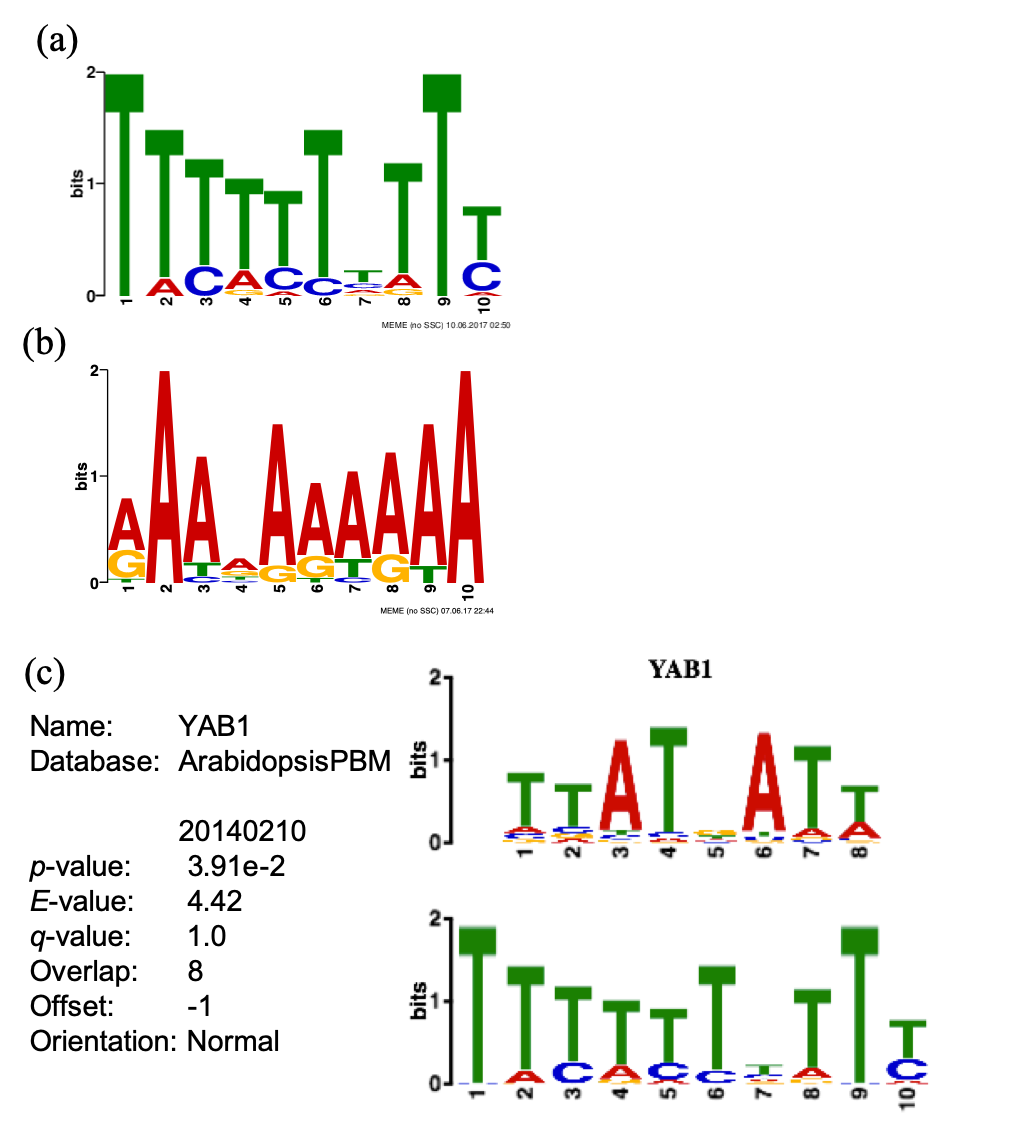

Figure S7. Significant motif present in *Micromonas pusilla* RCC299 gene sequences described in Table S4. The motif consensus logo for the positive, or coding strand (a), and for the negative strand (b) are shown. The best match for the consensus motif in the *Arabidopsis* regulatory element database (Franco-Zorilla *et al*., 2014) is to the element YAB1 (c).

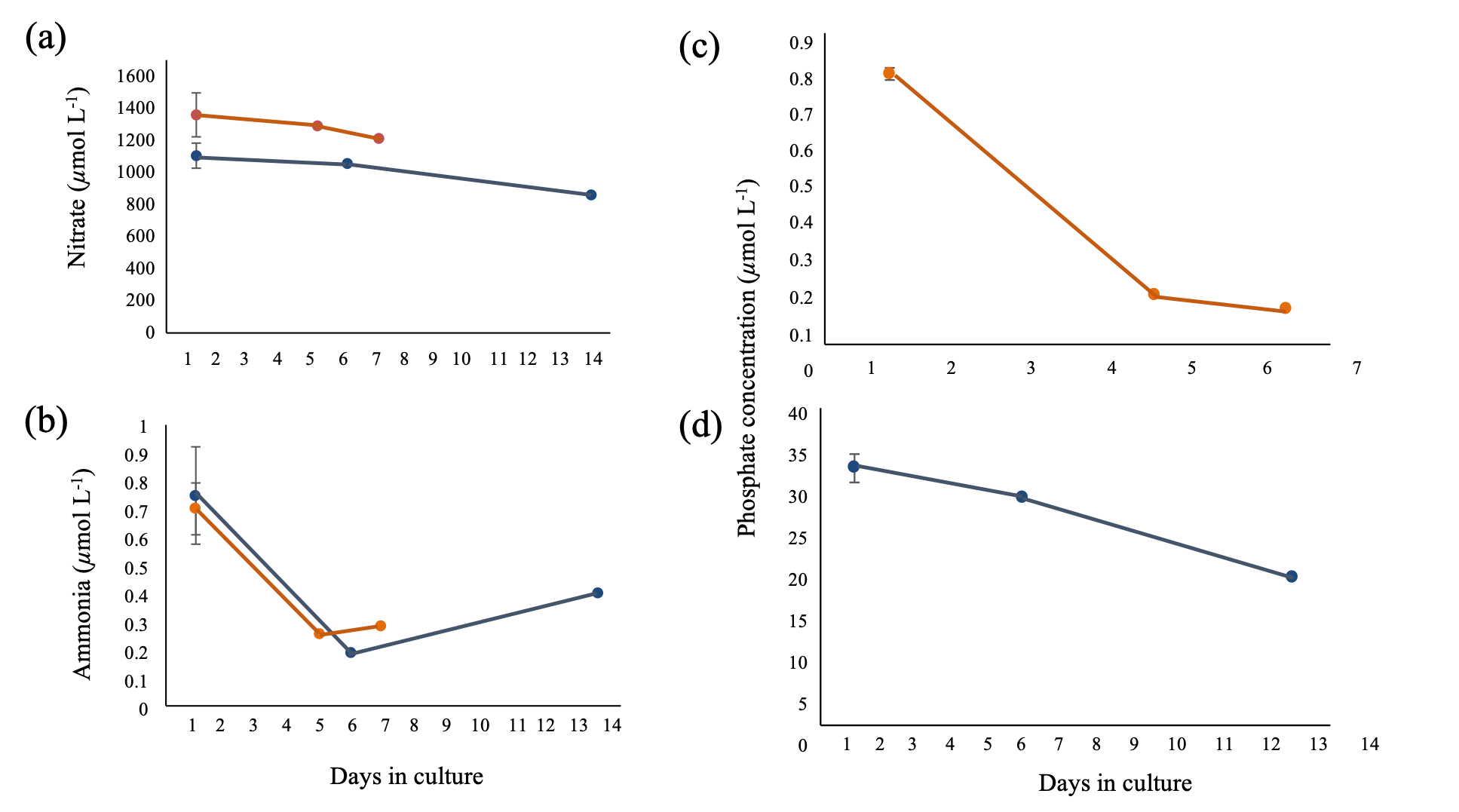

Figure S8. Nitrate, ammonium, and phosphate concentrations at each sampling point for *Micromonas pusilla* CCMP 1545 cultures. Average concentrations of nitrate (a), ammonia (b), phosphate in P-deficient (c), and phosphate in P-replete treatments (d) with standard deviation for P-replete (blue) and P-deficient (orange) treatments.

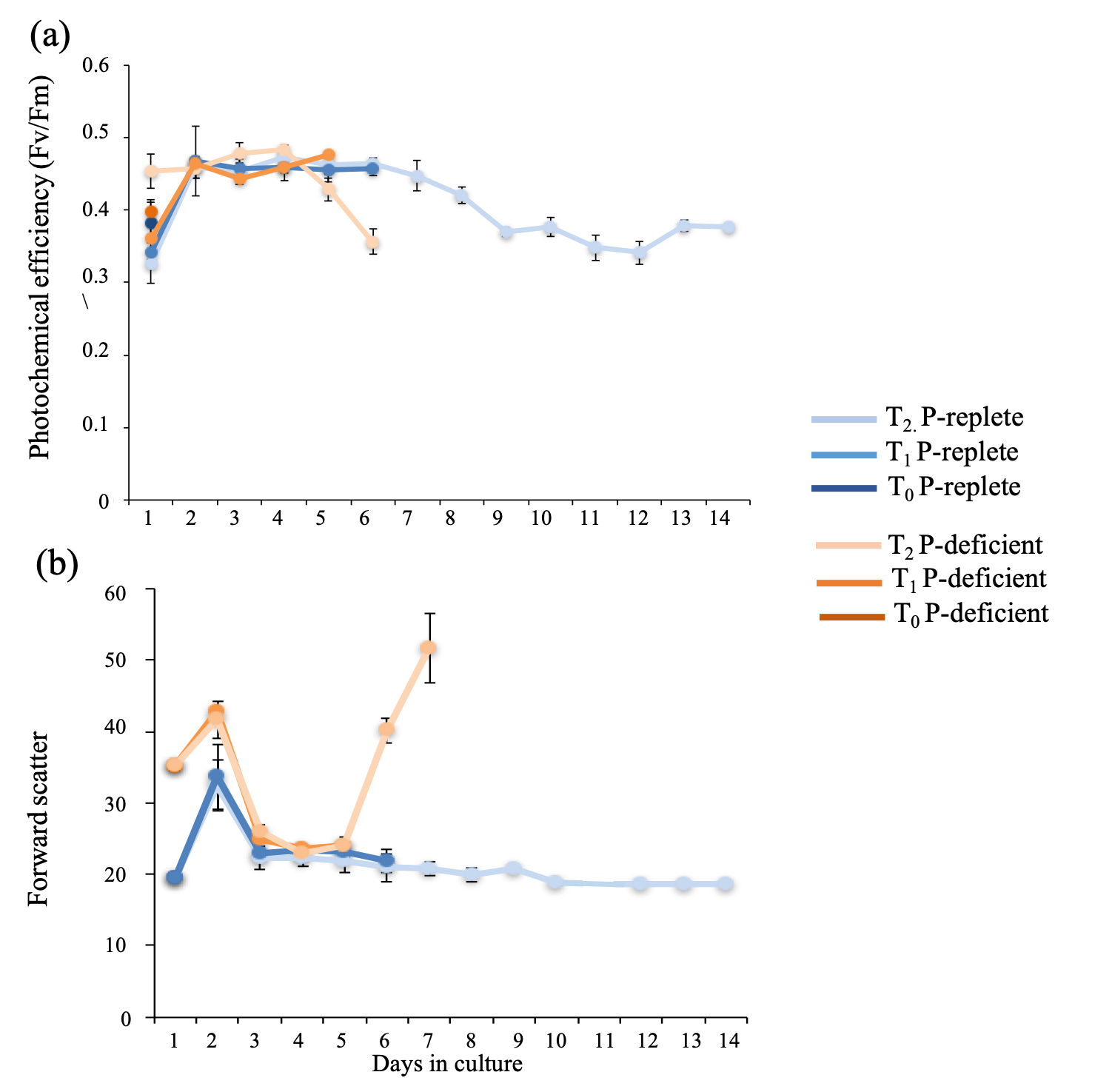

Figure S9. Photochemical efficiency and cell size during the experiment. Averages with standard deviation are shown for P-replete (blue) and P-deficient (orange) treatments. Photochemical efficiency (a) was measured at the same time each day using Fluorescence Induction and Relaxation (FIRe). Forward scatter of the cells in culture (b) was measured using flow cytometry and was used as a proxy for cell size. Forward scatter data was not collected on day 12.

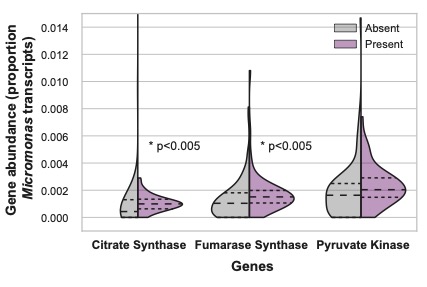

Figure S10. Normalized expression of three *Micromonas* carbon-cycle genes in the presence or absence of *Micromonas* *psr*1-like genes across the TARA Oceans dataset. The three genes, citrate synthase, fumarase synthase, and pyruvate kinase, all have a role in the tricarboxylic acid (TCA) cycle and were found to be differentially regulated under phosphorus-deficient conditions in *M. pusilla* and *M. commoda*. The normalized expression of these genes is plotted as a percentage of the total *Micromonas* transcript pool by sample. Violin plots represent the distribution of the normalized expression of the genes in the presence (purple) or absence (gray) of the *psr*1-like gene in the *Micromonas* transcriptome pool by sample.  The larger dashed line represents the distribution mean and the smaller dashed lines represent the 25% and 75% quartiles.  Asterisks indicate a significant difference in distribution of gene expression with and without the expression of *Micromonas psr*1-like genes (Kolmogorov-Smirnov, p < 0.005).

Table S1. Enzymes that contain a significant motif in the gene sequences of *M. pusilla* CCMP1545 (FrozenGeneCatalog_20080206 (ver 1)). Column 2: the Enzyme Commission (E.C.) number or pfam identifier; Column 3: main pathway(s) in which the enzyme is involved. These genes were used in the initial analysis of *M. pusilla* genes for a significant motif.

| **Enzyme** | **E.C. Number** | **Pathway** |
| --- | --- | --- |
| Glyceraldehyde-3-phosphate dehydrogenase | 1.2.1.12 | Calvin cycle (Chloroplast) |
| Glyceraldehyde-3-phosphate dehydrogenase | 1.2.1.12 | Glycolysis |
| Phosphoglycerate kinase | 2.7.2.3 | Glycolysis |
| Pyruvate kinase | 2.7.1.40 | Glycolysis |
| Citrate synthase | 2.3.3.1 | TCA^a^ cycle |
| Malate dehydrogenase | 1.1.1.37 | TCA cycle |
| Succinate dehydrogenase | 1.3.5.1 | TCA cycle and ETC^b^ |
| Fumarase | 4.2.1.2 | TCA cycle |
| Proline oxidase | 1.5.1.2 | Arginine and proline metabolism |
| Aspartate transcarbamylase | 2.1.3.2 | Pyrimidine biosynthesis |
| NaPO_4_ transporter | PF02690 | inorganic nutrient transport |
| Long chain fatty acid CoA-ligase | 6.2.1.3 | Fatty acid metabolism |
| Diacylglycerol O-acetyltransferase | 2.3.1.20 | Glycerolipid, TAG^c^ biosynthesis |

^a^Tricarboxylic acid cycle

^b^Electron transport chain

^c^Triacylglycerol

Table S2. Significant matches to the *psr*1-like gene predicted protein sequence of *Micromonas pusilla* CCMP1545 from the marine microbial eukaryote transcriptome sequencing project (MMETSP). Bolded names indicate taxa with protein sequences that contain an N-terminal myb-like DNA-binding domain and a C-terminal myb coiled-coil domain, whereas non-bolded taxa contain only the DNA-binding domain.

| **SampleName** | **Phylum** | **Class** | **Genus** | **Species** | **Strain** |
| --- | --- | --- | --- | --- | --- |
| **MMETSP0063** | **Chlorophyta** | **Chlorophyceae** | **Chlamydomonas** | **euryale** | **CCMP219** |
| **MMETSP1391** | Chlorophyta | Chlorophyceae | Chlamydomonas | leiostraca | SAG11-49 |
| **MMETSP0052** | Chlorophyta | Chlorophyceae | Polytomella | parva | SAG63-3 |
| **MMETSP1127** | **Chlorophyta** | **Chlorophyceae** | **Dunaliella** | **tertiolecta** | **CCMP1320** |
| **MMETSP1128** | **Chlorophyta** | **Chlorophyceae** | **Dunaliella** | **tertiolecta** | **CCMP1320** |
| **MMETSP1180** | Chlorophyta | Chlorophyceae | Chlamydomonas | sp | CCMP681 |
| **MMETSP1159** | Chlorophyta | Chlorophytaincertaesedis | Picocystis | salinarum | CCMP1897 |
| **MMETSP1460** | **Chlorophyta** | **Mamiellophyceae** | **Bathycoccus** | **prasinos** | **RCC716** |
| **MMETSP1106** | **Chlorophyta** | **Mamiellophyceae** | **Mantoniella** | **antarctica** | **SL-175** |
| **MMETSP1468** | Chlorophyta | Mamiellophyceae | Mantoniella | sp. | CCMP1436 |
| **MMETSP1327** | **Chlorophyta** | **Mamiellophyceae** | **Micromonas** | **pusilla** | **RCC2306** |
| **MMETSP1326** | Chlorophyta | Mamiellophyceae | Unidentifiedeukaryote | sp. | RCC2288 |
| **MMETSP0419** | **Chlorophyta** | **Prasinophyceae** | **Tetraselmis** | **sp.** | **GSL018** |
| **MMETSP0491** | Chlorophyta | Prasinophyceae | Tetraselmis | chuii | PLY429 |
| **MMETSP0804** | **Chlorophyta** | **Prasinophyceae** | **Tetraselmis** | **astigmatica** | **CCMP880** |
| **MMETSP0817** | Chlorophyta | Prasinophyceae | Tetraselmis | striata | LANL1001 |
| **MMETSP0818** | Chlorophyta | Prasinophyceae | Tetraselmis | striata | LANL1001 |
| **MMETSP0819** | Chlorophyta | Prasinophyceae | Tetraselmis | striata | LANL1001 |
| **MMETSP0820** | Chlorophyta | Prasinophyceae | Tetraselmis | striata | LANL1001 |
| **MMETSP1438** | Chlorophyta | Prasinophyceae | Pterosperma | sp. | CCMP1384 |
| **MMETSP1399** | **Chlorophyta** | **Prasinophyceae** | **Bathycoccus** | **prasinos** | **CCMP1898** |
| **MMETSP0033** | Chlorophyta | Prasinophyceae | Dolichomastix | tenuilepis | CCMP3274 |
| **MMETSP1387** | **Chlorophyta** | **Prasinophyceae** | **Micromonas** | **sp.** | **RCC472** |

Table S2. Continued.

| **SampleName** | **Phylum** | **Class** | **Genus** | **Species** | **Strain** |
| --- | --- | --- | --- | --- | --- |
| **MMETSP1393** | **Chlorophyta** | **Prasinophyceae** | **Micromonas** | **sp.** | **CS-222** |
| **MMETSP1400** | **Chlorophyta** | **Prasinophyceae** | **Micromonas** | **sp.** | **RCC451** |
| **MMETSP1401** | **Chlorophyta** | **Prasinophyceae** | **Micromonas** | **pusilla** | **CCAC1681** |
| **MMETSP1402** | **Chlorophyta** | **Prasinophyceae** | **Micromonas** | **pusilla** | **RCC1614** |
| **MMETSP1403** | **Chlorophyta** | **Prasinophyceae** | **Micromonas** | **pusilla** | **CCMP1723** |
| **MMETSP1404** | **Chlorophyta** | **Prasinophyceae** | **Micromonas** | **pusilla** | **CCMP494** |
| **MMETSP0929** | **Chlorophyta** | **Prasinophyceae** | **Ostreococcus** | **mediterraneus** | **clade-D-RCC2572** |
| **MMETSP0930** | **Chlorophyta** | **Prasinophyceae** | **Ostreococcus** | **mediterraneus** | **clade-D-RCC1621** |
| **MMETSP0938** | **Chlorophyta** | **Prasinophyceae** | **Ostreococcus** | **mediterraneus** | **clade-D-RCC1107** |
| **MMETSP0939** | **Chlorophyta** | **Prasinophyceae** | **Ostreococcus** | **lucimarinus** | **clade-A-BCC118000** |
| **MMETSP0803** | Chlorophyta | Prasinophyceae | Crustomastix | stigmata | CCMP3273 |
| **MMETSP0034** | Chlorophyta | Prasinophyceae | Nephroselmis | pyriformis | CCMP717 |
| **MMETSP1315** | Chlorophyta | Prasinophyceae | Prasinoderma | singularis | RCC927 |
| **MMETSP1316** | **Chlorophyta** | **Prasinophyceae** | **Pycnococcus** | **provasolii** | **RCC2336** |
| **MMETSP1459** | Chlorophyta | Prasinophyceae | Pycnococcus | provasolii | RCC931 |
| **MMETSP1471** | Chlorophyta | Prasinophyceae | Pycnococcus | provasolii | RCC733 |
| **MMETSP1472** | Chlorophyta | Prasinophyceae | Pycnococcus | provasolii | RCC251 |
| **MMETSP0941** | **Chlorophyta** | **Prasinophyceae** | **Prasinococcus** | **capsulatus** | **CCMP1194** |
| **MMETSP0806** | Chlorophyta | Prasinophyceae | Prasinoderma | coloniale | CCMP1413 |
| **MMETSP1085** | Chlorophyta | Prasinophyceae | Pycnococcus | sp | CCMP1998 |
| **MMETSP0058** | **Chlorophyta** | **Prasinophyceae** | **Pyramimonas** | **parkeae** | **CCMP726** |
| **MMETSP0059** | **Chlorophyta** | **Prasinophyceae** | **Pyramimonas** | **parkeae** | **CCMP726** |
| **MMETSP1445** | **Chlorophyta** | **Prasinophyceae** | **Pyramimonas** | **sp.** | **CCMP2087** |

Table S2. Continued.

| **SampleName** | **Phylum** | **Class** | **Genus** | **Species** | **Strain** |
| --- | --- | --- | --- | --- | --- |
| **MMETSP1169** | **Chlorophyta** | **Prasinophyceae** | **Pyramimonas** | **obovata** | **CCMP722** |
| **MMETSP1473** | Chlorophyta | Trebouxiophyceae | Stichococcus | sp. | RCC1054 |
| **MMETSP1330** | **Chlorophyta** | **Trebouxiophyceae** | **Picochlorum** | **sp.** | **RCC944** |
| **MMETSP0807** | **Chlorophyta** | **Unknown** | **Picocystis** | **salinarum** | **CCMP1897** |
| **MMETSP0126** | **Ciliophora** | **Spirotrichea** | **Strombidinopsis** | **acuminatum** | **SPMC142** |
| **MMETSP0123** | Ciliophora | Spirotrichea | Favella | ehrenbergii | Fehren1 |
| **MMETSP0384** | Dinophyta | Dinophyceae | Alexandrium | tamarense | CCMP1771 |
| **MMETSP0470** | Dinophyta | Dinophyceae | Oxyrrhis | marina |  |
| **MMETSP1367** | Dinophyta | Dinophyceae | Symbiodinium | sp. | C1 |
| **MMETSP1369** | Dinophyta | Dinophyceae | Symbiodinium | sp. | C1 |
| **MMETSP1086** | Glaucophyta | Glaucocystophyceae | Cyanoptyche | gloeocystis | SAG4.97 |
| **MMETSP0308** | Glaucophyta | Glaucophyceae | Gloeochaete | wittrockiana | SAG46.84 |
| **MMETSP1089** | Glaucophyta | Glaucophyceae | Gloeochaete | wittrockiana | SAG46.84 |
| **MMETSP1333** | **Haptophyta** | **Coccolithophyceae** | **Scyphosphaera** | **apsteinii** | **RCC1455** |
| **MMETSP1464** | Haptophyta | Haptophyceae | Exanthemachrysis | gayraliae | RCC1523 |
| **MMETSP1139** | Haptophyta | Pavlovophyceae | Pavlova | sp. | CCMP459 |
| **MMETSP1140** | Haptophyta | Pavlovophyceae | Pavlova | sp. | CCMP459 |
| **MMETSP1381** | Haptophyta | Pavlovophyceae | Pavlova | sp. | CCMP459 |
| **MMETSP1463** | Haptophyta | Pavlovophyceae | Pavlova | lutheri | RCC1537 |
| **MMETSP1466** | Haptophyta | Pavlovophyceae | Pavlova | gyrans | CCMP608 |
| **MMETSP1334** | **Haptophyta** | **Prymnesiophyceae** | **Calcidiscus** | **leptoporus** | **RCC1130** |
| **MMETSP0164** | Haptophyta | Prymnesiophyceae | Coccolithus | pelagicusssp.braarudi | PLY182g |
| **MMETSP1136** | Haptophyta | Prymnesiophyceae | Pleurochrysis | carterae | CCMP645 |
| **MMETSP1138** | Haptophyta | Prymnesiophyceae | Pleurochrysis | carterae | CCMP645 |
| **MMETSP0595** | **Haptophyta** | **Prymnesiophyceae** | **Isochrysis** | **galbana** | **CCMP1323** |

Table S2. Continued.

| **SampleName** | **Phylum** | **Class** | **Genus** | **Species** | **Strain** |
| --- | --- | --- | --- | --- | --- |
| **MMETSP0943** | Haptophyta | Prymnesiophyceae | Isochrysis | galbana | CCMP1323 |
| **MMETSP0944** | Haptophyta | Prymnesiophyceae | Isochrysis | galbana | CCMP1323 |
| **MMETSP1090** | Haptophyta | Prymnesiophyceae | Isochrysis | sp. | CCMP1244 |
| **MMETSP1129** | **Haptophyta** | **Prymnesiophyceae** | **Isochrysis** | **sp.** | **CCMP1324** |
| **MMETSP1131** | Haptophyta | Prymnesiophyceae | Isochrysis | sp. | CCMP1324 |
| **MMETSP1132** | Haptophyta | Prymnesiophyceae | Isochrysis | sp. | CCMP1324 |
| **MMETSP1388** | Haptophyta | Prymnesiophyceae | Isochrysis | sp. | CCMP1244 |
| **MMETSP0994** | Haptophyta | Prymnesiophyceae | Emiliania | huxleyi | 379 |
| **MMETSP0995** | Haptophyta | Prymnesiophyceae | Emiliania | huxleyi | 379 |
| **MMETSP0996** | Haptophyta | Prymnesiophyceae | Emiliania | huxleyi | 379 |
| **MMETSP0997** | Haptophyta | Prymnesiophyceae | Emiliania | huxleyi | 379 |
| **MMETSP1006** | Haptophyta | Prymnesiophyceae | Emiliania | huxleyi | 374 |
| **MMETSP1007** | Haptophyta | Prymnesiophyceae | Emiliania | huxleyi | 374 |
| **MMETSP1008** | Haptophyta | Prymnesiophyceae | Emiliania | huxleyi | 374 |
| **MMETSP1009** | Haptophyta | Prymnesiophyceae | Emiliania | huxleyi | 374 |
| **MMETSP1150** | Haptophyta | Prymnesiophyceae | Emiliania | huxleyi | PLYM219 |
| **MMETSP1151** | Haptophyta | Prymnesiophyceae | Emiliania | huxleyi | PLYM219 |
| **MMETSP1152** | Haptophyta | Prymnesiophyceae | Emiliania | huxleyi | PLYM219 |
| **MMETSP1153** | Haptophyta | Prymnesiophyceae | Emiliania | huxleyi | PLYM219 |
| **MMETSP1154** | Haptophyta | Prymnesiophyceae | Emiliania | huxleyi | CCMP370 |
| **MMETSP1155** | Haptophyta | Prymnesiophyceae | Emiliania | huxleyi | CCMP370 |
| **MMETSP1156** | Haptophyta | Prymnesiophyceae | Emiliania | huxleyi | CCMP370 |
| **MMETSP1157** | Haptophyta | Prymnesiophyceae | Emiliania | huxleyi | CCMP370 |
| **MMETSP1363** | Haptophyta | Prymnesiophyceae | Gephyrocapsa | oceanica | RCC1303 |

Table S2. Continued.

| **SampleName** | **Phylum** | **Class** | **Genus** | **Species** | **Strain** |
| --- | --- | --- | --- | --- | --- |
| **MMETSP1364** | Haptophyta | Prymnesiophyceae | Gephyrocapsa | oceanica | RCC1303 |
| **MMETSP1365** | **Haptophyta** | **Prymnesiophyceae** | **Gephyrocapsa** | **oceanica** | **RCC1303** |
| **MMETSP1366** | **Haptophyta** | **Prymnesiophyceae** | **Gephyrocapsa** | **oceanica** | **RCC1303** |
| **MMETSP1100** | Haptophyta | Prymnesiophyceae | Phaeocystis | antarctica | CaronLabIsolate |
| **MMETSP1444** | Haptophyta | Prymnesiophyceae | Phaeocystis | antarctica | CCMP1374 |
| **MMETSP1465** | **Haptophyta** | **Prymnesiophyceae** | **Phaeocystis** | **cordata** | **RCC1383** |
| **MMETSP1162** | Haptophyta | Prymnesiophyceae | Phaeocystis | sp. | CCMP2710 |
| **MMETSP0006** | **Haptophyta** | **Prymnesiophyceae** | **Prymnesium** | **parvum** | **Texoma1** |
| **MMETSP0007** | Haptophyta | Prymnesiophyceae | Prymnesium | parvum | Texoma1 |
| **MMETSP0008** | Haptophyta | Prymnesiophyceae | Prymnesium | parvum | Texoma1 |
| **MMETSP0008** | Haptophyta | Prymnesiophyceae | Prymnesium | parvum | Texoma1 |
| **MMETSP0814** | **Haptophyta** | **Prymnesiophyceae** | **Prymnesium** | **parvum** | **Texoma1** |
| **MMETSP0815** | **Haptophyta** | **Prymnesiophyceae** | **Prymnesium** | **parvum** | **Texoma1** |
| **MMETSP1083** | **Haptophyta** | **Prymnesiophyceae** | **Prymnesium** | **parvum** | **Texoma1** |
| **MMETSP0143** | **Haptophyta** | **Prymnesiophyceae** | **Chrysochromulina** | **polylepis** | **CCMP1757** |
| **MMETSP0145** | Haptophyta | Prymnesiophyceae | Chrysochromulina | polylepis | CCMP1757 |
| **MMETSP0146** | Haptophyta | Prymnesiophyceae | Chrysochromulina | polylepis | CCMP1757 |
| **MMETSP0147** | **Haptophyta** | **Prymnesiophyceae** | **Chrysochromulina** | **polylepis** | **CCMP1757** |
| **MMETSP0286** | **Haptophyta** | **Prymnesiophyceae** | **Chrysochromulina** | **polylepis** | **UIO037** |
| **MMETSP0287** | **Haptophyta** | **Prymnesiophyceae** | **Chrysochromulina** | **rotalis** | **UIO044** |
| **MMETSP1094** | **Haptophyta** | **Prymnesiophyceae** | **Chrysochromulina** | **brevifilum** | **UTEXLB985** |
| **MMETSP1096** | **Haptophyta** | **Prymnesiophyceae** | **Chrysochromulina** | **ericina** | **CCMP281** |
| **MMETSP1335** | Haptophyta | Prymnesiophyceae | Chrysoculter | rhomboideus | RCC1486 |
| **MMETSP1474** | **Haptophyta** | **Prymnesiophyceae** | **Imantonia** | **sp.** | **RCC918** |

Table S2. Continued.

| **SampleName** | **Phylum** | **Class** | **Genus** | **Species** | **Strain** |
| --- | --- | --- | --- | --- | --- |
| **MMETSP1178** | Haptophyta | Prymnesiophyceae | Unidentifiedeukaryote | sp. | CCMP2000 |
| **MMETSP0312** | Rhodophyta | Compsopogonophyceae | Compsopogon | coeruleus | SAG36.94 |
| **MMETSP1172** | Rhodophyta | Porphyridiophyceae | Timspurckia | oligopyrenoides | CCMP3278 |
| **MMETSP0313** | Rhodophyta | Rhodellophyceae | Porphyridium | aerugineum | SAG1380-2 |
| **MMETSP0167** | Rhodophyta | Rhodellophyceae | Rhodella | maculata | CCMP736 |
| **MMETSP0314** | Rhodophyta | Rhodellophyceae | Rhodella | maculata | CCMP736 |
| **MMETSP0011** | Rhodophyta | Rhodellophyceae | Rhodosorus | marinus | CCMP769 |
| **MMETSP0315** | Rhodophyta | Rhodellophyceae | Rhodosorus | marinus | UTEXLB2760 |
| **MMETSP0417** | Unknown | Lobosa | Mayorella | sp | BSH-02190019 |
| **MMETSP0780** | Unknown | Unknown | Palpitomonas | bilix | NIES-2562 |
| **MMETSP0447** | Unknown | Unknown | Stygamoeba | regulata | BSH-02190019 |
| **MMETSP0098** | Unknown | Unknown | Unidentifiedeukaryote | sp. | NY0313808BC1 |
| **MMETSP0099** | Unknown | Unknown | Unidentifiedeukaryote | sp. | NY0313808BC1 |
| **MMETSP0100** | Unknown | Unknown | Unidentifiedeukaryote | sp. | NY0313808BC1 |
| **MMETSP0982** | Unknown | Unknown | Unidentifiedeukaryote | sp. | CCMP2436 |
| **MMETSP0983** | Unknown | Unknown | Unidentifiedeukaryote | sp. | CCMP2436 |
| **MMETSP0984** | Unknown | Unknown | Unidentifiedeukaryote | sp. | CCMP2436 |
| **MMETSP0985** | Unknown | Unknown | Unidentifiedeukaryote | sp. | CCMP2436 |
| **MMETSP1084** | **Unknown** | **Unknown** | **Micromonas** | **sp.** | **RCC472** |
| **MMETSP1309** | Unknown | Unknown | Unidentifiedeukaryote | sp. | RCC998 |
| **MMETSP1310** | **Unknown** | **Unknown** | **Unidentifiedeukaryote** | **sp.** | **RC2339** |
| **MMETSP1311** | **Unknown** | **Unknown** | **Unidentifiedeukaryote** | **sp.** | **RCC856** |

Table S2. Continued.

| **SampleName** | **Phylum** | **Class** | **Genus** | **Species** | **Strain** |
| --- | --- | --- | --- | --- | --- |
| **MMETSP1312** | **Unknown** | **Unknown** | **Unidentifiedeukaryote** | **sp.** | **RCC2335** |
| **MMETSP1433** | Unknown | Unknown | Unidentifiedeukaryote | sp. | NY070348D |
| **MMETSP1446** | **Unknown** | **Unknown** | **Unidentifiedeukaryote** | **sp.** | **CCMP2111** |
| **MMETSP1453** | **Unknown** | **Unknown** | **Unidentifiedeukaryote** | **sp.** | **RCC701** |
| **MMETSP1456** | **Unknown** | **Unknown** | **Unidentifiedeukaryote** | **sp.** | **RCC1871** |
| **MMETSP1469** | Unknown | Unknown | Unidentifiedeukaryote | sp. | CCMP1205 |
| **MMETSP1470** | **Unknown** | **Unknown** | **Unidentifiedeukaryote** | **sp.** | **CCMP2175** |

Table S3. Significant matches to the motif discovered in *Micromonas pusilla* CCMP1545 genes. The conserved motif was searched against the *Arabidopsis* *thaliana* PBM motifs database (Franco-Zorilla *et al*., 2014). The consensus motif and significant matches based on the two *M. pusilla* CCMP1545 gene models available in the Joint Genome Institute database are shown.

| **Consensus** | **Gene model** | **Match** | ***p*-value** | **E-value** | **q-value** |
| --- | --- | --- | --- | --- | --- |
| GCGTGGTGAA | MBARI_models (ver 1) | REM1_2 | 0.007 | 0.86 | 1 |
|  |  | AT5G28300 | 0.028 | 3.22 | 1 |
|  |  | MYB111_2 | 0.043 | 4.88 | 1 |
|  |  | ANAC46 | 0.049 | 5.56 | 1 |
| AAACAAACGA | FrozenGeneCatalog_20080206 (ver 1) (archived) | TOE1_2 | 0.009 | 1.06 | 1 |
|  |  | TOE2_2 | 0.01 | 1.15 | 1 |
|  |  | ARR11 | 0.025 | 2.82 | 1 |
|  |  | MYB52 | 0.026 | 3 | 1 |
|  |  | MYB55 | 0.053 | 5.98 | 1 |

Table S4. Genes from *Micromonas commoda* RCC299 that were differentially expressed under phosphorus deficiency (Whitney & Lomas, 2016). Query of the dataset was focused on genes related to the present study (see methods in main text). The Joint Genome Institute protein identification number, the name of the enzyme that the gene encodes, fold change in gene expression, and the pathway in which the enzyme is involved are listed. Fold change data are from Whitney & Lomas (2016). Bolded protein IDs and enzyme names indicate these gene sequences contained the motif shown in Figure S7.

| **Protein ID** | **Enzyme** | **P*i*-deficient (log2 fold change)** | **Pathway** |
| --- | --- | --- | --- |
| 60184 | Myb family transcription factor (putative psr1-like gene) | 4.3 | Transcription factor |
| **75917** | **Malate dehydrogenase** | 1.08 | TCA |
| **96682** | **Citrate synthase** | -1.65 | TCA |
| **96474** | **Fumarase** | 1.09 | TCA |
| **96455** | **Pyruvate carboxylase** | 2.15 | TCA |
| **100385** | **Proline oxidase** | 1.10 | Amino acid metabolism |
| 99771 | Aspartate aminotransferase | 1.25 | Amino acid metabolism |
| 115504.8 | D-aspartate oxidase | 1.18 | Amino acid metabolism and anaplerotic pathways |
| **123909.7** | **Asparagine synthase (glutamine-hydrolysing)** | 1.15 | Amino acid metabolism |
| **56785** | **Acetate--CoA ligase** | 1.69 | Glycolysis |
| **104803** | **Phosphoglycerate kinase*** | 1.69 | Glycolysis |
| 60369 | Phosphoglycerate kinase | 2.15 | Glycolysis |
| **82586** | **Pyruvate kinase*** | 4.53 | Glycolysis |
| 38152 | Pyruvate kinase | 1.87 | Glycolysis |
| **104954** | **Glyceraldehyde 3-phosphate dehydrogenase** | 4.03 | Carbon fixation |
| 104767 | Phosphoglycerate kinase | -1.45 | Carbon fixation |

Table S4. Continued.

| **Protein ID** | **Enzyme** | **P*i*-deficient (log2 fold change)** | **Pathway** |
| --- | --- | --- | --- |
| **106294** | **5'-nucleotidase** | 1.4 | Nucleotide metabolism |
| **86041** | **Triosephosphate isomerase** | -0.95 | Carbon fixation |
| **61667** | **Glutamate-1-semialdehyde 2,1-aminomutase** | 1.23 | Carbon fixation |
| 109214 | triose phosphate:phosphate translocator | 3.89 | Export triose sugar |
| **97893** | **Biotin synthase** | 2.29 | Vitamin metabolism |
| **90156** | **Dihydroxy-acid dehydratase** | 1.95 | Pantothenate and CoA biosynthesis |
| 107432 | 3-oxoacyl-[acyl-carrier protein] synthase | 0.77 | Fatty acid biosynthesis |
| 109209 | 3-oxoacyl-[acyl-carrier protein] synthase | 1.02 | Fatty acid biosynthesis |
| 85187 | 3-oxoacyl-[acyl-carrier protein] reductase | -1.41 | Fatty acid biosynthesis |
| 94289 | 1-acylglycerol-3-phosphate O-acyltransferase | -1.01 | Glycerolipid metabolism |
| **58855** | **Phosphatidic acid phosphatase** | 2.10 | Glycerolipid metabolism |
| **78992** | **Lysophospholipid acyltransferase** | 1.56 | Glycerolipid metabolism |
| 80225 | Acylglycerol lipase | 1.40 | Glycerolipid metabolism |
| 79438 | Glycerol kinase | -1.00 | Glycerolipid metabolism |
| **80077** | **dTDP-glucose 4,6-dehydratase** | 2.87 | Secondary metabolite biosynthesis |

Table S5. Characteristics and comparisons of the *psr*1 gene in model organisms and in *Micromonas*. Coverage (covg), percent identity, and the E-value are shown for each comparison. Comparisons are based on pair-wise alignments of the amino acid-derived sequences using NCBI.

|  | Length | *C. reinhardtii* cc 125 | | | *M. commoda* RCC299 | | | |
| --- | --- | --- | --- | --- | --- | --- | --- | --- |
|  | nt (aa) | Covg (%) | Identity (%) | E-value | | Covg (%) | Identity (%) | E-value |
| *C. reinhardtii (*cc125) | 4,052 (752) |  |  |  | |  |  |  |
| *M. pusilla* (CCMP1545) | 1,284 (428) | 23 | 66 | 2 x 10^-24^ | | 68 | 47 | 5 x 10^-45^ |
| *M. commoda* (RCC299) | 1,486 (466) | 22 | 59 | 1 x 10^-26^ | |  |  |  |
| *A. thaliana* | 2482 (409) | 37 | 70 | 3 x 10^-27^ | |  |  |  |

nt = nucleotide

aa = amino acid
